# Evolverator: An engineered yeast system to drive rapid continuous *in vivo* evolution of proteins

**DOI:** 10.1101/2024.09.01.610536

**Authors:** Asli Azizoglu, Eline Y. Bijman, Roger Brent

**Author notes:** Equal contribution (Asli Azizoglu, Eline Y. Bijman). Corresponding author. A.A.

## Abstract

Continuous *in vivo* evolution of non-essential proteins requires an engineered link between target protein and host cell fitness, which is difficult to establish and vulnerable to escape mutations. We systematically designed and tested *Saccharomyces cerevisiae* gene circuits guided by quantitative mathematical models that predicted circuit, organism, and population behavior. We identified four generalizable design principles for escape-resistant continuous evolution and built the Evolverator, a yeast platform that couples protein binding to proliferation rate while suppressing escape mutations. We used the Evolverator to improve human estrogen receptor ligand affinity and generate stronger yeast repressors from the bacterial LacI protein. The resulting variants revealed previously undescribed hER substitutions within and outside the binding pocket, as well as LacI mutations that enhance repression in yeast despite reduced activity in *Escherichia coli*. Our work establishes a generalizable, model-guided framework for building yeast strains to allow continuous evolution of diverse protein functions *in vivo*.

## INTRODUCTION

Continuous *in vivo* directed evolution couples continuous mutagenesis of a target protein inside the cell to fitness-based selection by linking the protein’s function to cell fitness. By applying gentle selective pressure that slowly and continuously increases over time, continuous evolution allows evolutionary trajectories that pass through transiently neutral or deleterious intermediates and generate variants that satifsfy an investigator-determined fitness goal. In other words, if a mutation does not show improvement, it is neither enriched nor is it immediately discarded. Compared to *in vitro* mutagenesis and non-fitness based selection methods, this approach widens the search space and phenotypes located across large fitness valleys can be accessed.

While continuous evolution used to be limited due to low *in vivo* mutagenesis rates, new methods such as base editors and error-prone replication like Orthorep [1], [2] have rates comparable to previously used *in vitro* methods[3]. However, continuous evolution also requires a link between the desired function of the evolving protein (phenotype) and cell fitness, and this is not always straightforward to establish. In most examples, this phenotype-fitness link is innate to the protein being evolved. For example, antibiotic resistance genes or genes encoding for essential enzymes can be easily evolved, as these are required for cell survival[2], [4]. However, for proteins without such a direct link, one must engineer it. A significant hurdle to this is escape mutations (also called cheater mutations) [5]. These are mutations that allow the cell to escape the engineered phenotype-fitness link and gain fitness without incorporating any beneficial mutations into the target protein. These must be prevented for successful continuous evolution.

The continuous evolution landscape has greatly expanded in the last decade. Phage-assisted continuous evolution (PACE) and its variants have moved beyond proof-of-concept to produce gene-therapy-relevant tools[6]. OrthoRep, initially a single-lab tool, has been adopted by external groups for applications and a variant called OptoRep that uses light to tune selection stringency demonstrated neofunctionalization of a yeast monocarboxylate transporter into a mevalonate importer [7], [8]. Mammalian continuous evolution has arrived through two independent routes: chimeric virus-like vesicles (PROTEUS[9]) and orthogonal RNA replication (REPLACE[10]). Most recently, Optovolution demonstrated light-directed evolution of a transcription factor in yeast by coupling its function to S-phase entry[11].

However, none of these systems provide a generalizable gene-circuit framework for engineering escape-resistant phenotype-fitness links for diverse non-essential protein functions in standard liquid culture, without specialized optogenetic hardware or continuous-flow equipment. Each existing platform suppresses escape mutations through system-specific design features rather than articulating transferable principles that a researcher could apply when building a new evolution system from scratch.

Here we describe the Evolverator, a generic, tunable selection system in *S. cerevisiae* that links target protein’s function (e.g., binding to a molecule) to cell fitness, while a built-in mutator continuously diversifies that protein until its function improves. Modular engineered gene circuits allow Evolverator cells to evolve the target protein toward experimenter-defined goals in long term, high throughput, liquid culture evolution campaigns, while escape mutations are being minimized. We demonstrate this using two proteins without any innate links to yeast cell fitness and meanwhile develop generalizable principles for escape mutation minimization. First, we evolve the human estrogen receptor ligand binding domain (hERLBD) to display higher affinities towards three different estrogen derivatives. In a second campaign, we evolve *E. coli lac* transcriptional repressor, LacI, toward increased transcriptional repression of an engineered yeast promoter. Additionally, we develop a parametrized model that identifies, before a strain is built, which circuit modifications (e.g. promoter strength, copy number, module stoichiometry) bring the expected affinity range and expression levels of a new target into the functional parameter space. Thus, providing a model-guided framework for a quantitative understanding of the system and for adapting the platform to other protein classes and functions. Overall, we engineered the most complex phenotype-fitness link for continuous evolution in a eukaryotic organism thus far, established generalizable principles for escape-resistant continuous evolution of non-essential eukaryotic proteins in high-throughput liquid culture, and demonstrated those principles through evolution campaigns in yeast.

## RESULTS

### Evolverator design to establish a phenotype-fitness link and automatic, tunable mutagenesis

We aimed to create a generalizable framework for continuous evolution of protein-ligand interactions. To this end, we sought a strategy in which weak protein-ligand binding would lead to (i) reduced fitness and (ii) increased mutagenesis of the target protein. This can be achieved by engineering cells such that the non-essential protein-ligand interaction controls both the expression of a mutagenesis protein and an essential gene in yeast. We designed a genetic circuit for this purpose that contains four components: (i) The Target protein, which when bound to its ligand, acts as a transcriptional repressor; (ii) the essential Growth protein; (iii) the Inverter protein, which is a repressor itself that converts the repressive action of the Target into expression of the Growth protein; and (iv) the Mutator protein, whose expression is repressed directly by the Target (Figure 1, Design). Consequently, cells carrying a low affinity Target should proliferate slowly and exhibit high Target mutagenesis rates, while cells with increased Target affinity should have reduced mutagenesis and proliferate more rapidly, allowing them to take over in liquid culture (Figure 1, Population Behavior).

**Figure 1.**
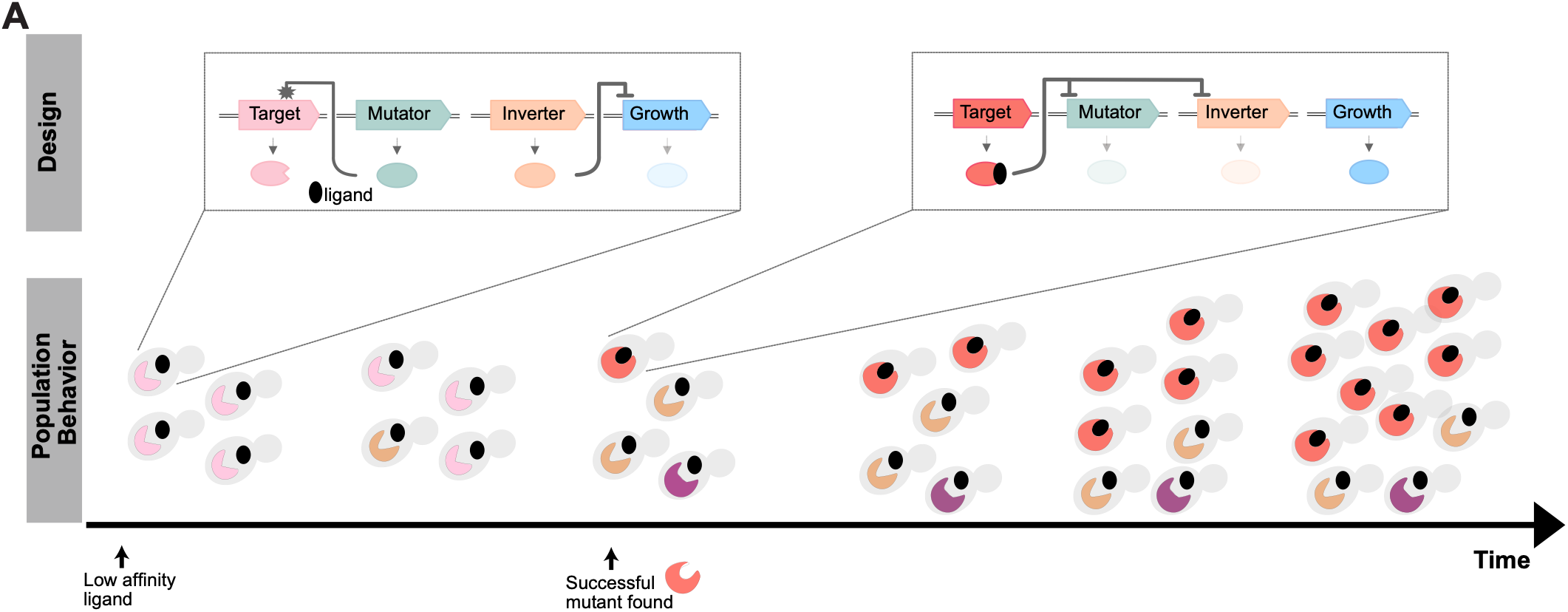
Overview of the Evolverator design. Design depicts the interactions between the molecular components. Population Behavior depicts growth behavior of cells with wild type receptor (light pink) and mutants (different colors) with different affinities towards the small molecule ligand. Cells that carry a higher affinity receptor proliferate faster.

Choosing the right essential gene for fitness control is critical for successful long-term evolution. We avoided antibiotic resistance genes and enzymes required for essential metabolites, as these can perturb cell physiology and cause gene circuits to behave unpredictably (for example, amino-acid depletion can impair translation of circuit components). Instead, we chose the essential membrane H^+^-ATPase, Pma1. Pma1 is long-lived and preferentially retained in mother cells during cell division[12], [13], which keeps mothers healthy during evolution while preventing production of viable daughters unless evolution succeeds. We previously showed that placing Pma1 under a TetR-repressible promoter enables growth rates from zero to maximum growth[14]. We therefore chose the engineered repressor TetR-Tup1 as our Inverter. Importantly, TetR-Tup1 repression can be inactivated by anhydrotetracycline. This allows us to maintain normal growth outside of evolution campaigns and to tune selection strength by adjusting Pma1 expression during evolution.

Our first goal was to increase ligand affinity of the human estrogen receptor ligand binding domain (hERLBD). We designed yeast promoters repressible by the fusion protein between bacterial DNA binding protein LexA and hERLBD, when hERLBD is bound to its ligand [15]. We verified that LexA-hERLBD exhibited different affinities for different estrogen-related ligands (Figure S1A), such that these ligands would then give us different starting points for evolution campaigns[16].

For the Mutator subsystem we used fusions between cytidine or adenine deaminases and T7 RNA polymerase [17], [18]. T7 polymerase binds a T7 promoter (P_T7_) placed before the open reading frame of the Target, then carries the deaminases downstream, introducing mutations along the gene. We used dA-T7, which contains the adenine deaminase abe8e (A>G mutations[19]), and dC-T7 with cytidine deaminase hA3A, (C>T mutations[20]). We tested those mutators in cells that contained a LexA-P_T7_-*URA3* construct and estimated mutagenesis rates from fluctuation assays [21]. dC-T7 and dA-T7 individually induced around 129×10^−7^ and 17×10^−7^ mutations per genome per generation, a 293- and 39-fold increase above background (Figure S1B). In cells that expressed both processive mutators, the combined mutation rate was 425-fold above background.

Overall, this gave us the four components needed: (i) LexA-hERLBD as the Target; (ii) Pma1 as the essential Growth protein; (iii) TetR-Tup1 as the Inverter protein; and (iv) dC-T7 and dA-T7 as the Mutator proteins.

### Design principles for escape-resistant continuous evolution

Any engineered phenotype-fitness link that requires the cell to find rare beneficial mutations in the target gene is vulnerable to escape mutations that break the engineered connection between the target protein’s function and cell fitness. We reasoned that a generalizable framework for continuous evolution must therefore be based on explicit design principles for suppressing escape mutations, and that those principles, once identified, would apply to any evolution system in which non-essential protein function is coupled to growth. In the course of systematic construction and testing of successive Evolverator strains, we identified four such design principles: (i) growth control must be separated from metabolic burden and established through asymmetrically inherited proteins, so that the link to fitness does not itself perturb the cell physiology it depends on; (ii) the key circuit components must be genetically redundant, so that loss-of-function mutations in any single copy cannot abolish the link; (iii) mutations that truncate the target protein must be selected against by coupling them to loss of an essential function; and (iv) endogenous genes that can substitute for the engineered growth-control gene must be identified and deleted. We describe the experiments that led to each of these principles below.

We combined LexA-hERLBD, TetR-Tup1, Pma1 and our mutators in one strain (Y3883) and grew 92 parallel cultures without β-estradiol (Round 1) for one growth cycle. As expected, this strain grew very slowly (Figure S2A), but between 50-80 hours, all cultures exhibited sharply increased growth rates (Figure 2A, 1x, and Figure S2B), indicating that each well had produced at least one cell that escaped *PMA1*-restricted growth control. This result demonstrated that the engineered connection between target protein function and cell proliferation was functional but that the single-copy inverter was the weak point, suggesting the second design principle: genetic redundancy in key circuit components (Figure 2B).

**Figure 2.**
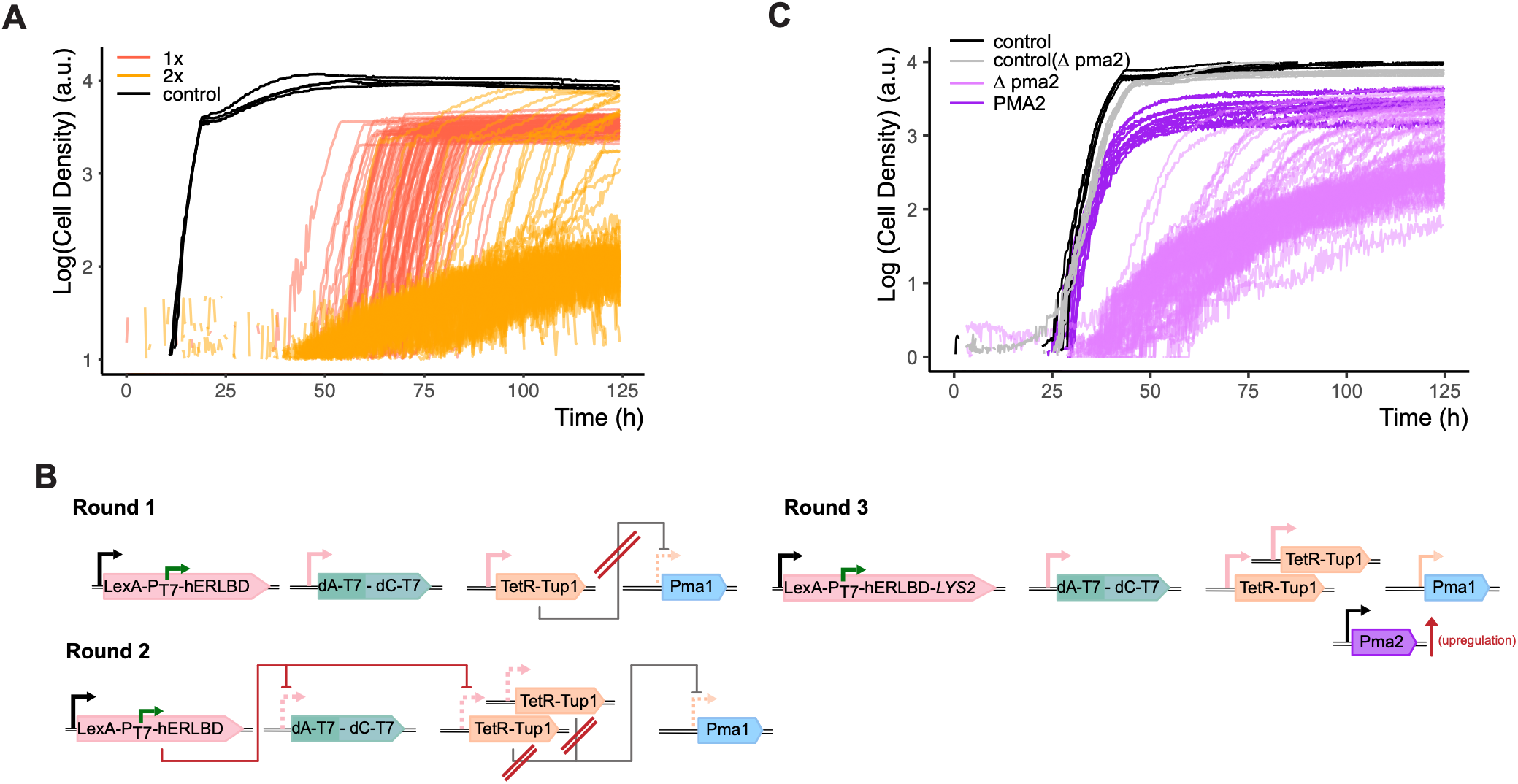
Design principles to prevent escape mutations. **A)** Growth of Y3883 with one genomic TetR-Tup1 copy (1x) and Y3884 with two genomic TetR-Tup1 copies (2x) in YPD without hormone, n=92 wells. Control: wells grown with 20 ng/mL aTc, n=4.**B)** Schematic depiction of the molecular components of cells used in each Round and the potential escape mutations in dark red. Active (solid) and inactive (dashed) promoters are the same color as the proteins that can repress them. Round 1: Y3883, Round 2: Y3884, Round 3: Y3740. **C)** Growth of Y3751 (*Δpma2*) and Y3740 (*PMA2)* as in (A), but with 10ng/mL aTc. Control wells grown with 20ng/mL aTc and 10*µ*M β-estradiol.

We generated a strain with a second TetR-Tup1 copy (Y3884) (Round 2) and grew it with low concentration of aTc (10ng/mL) to allow slow growth in the absence of β-estradiol (Figure S2C). In this round, only 20 out of 92 wells showed a sudden increase in growth rate indicative of escape mutations (Figure 2A, 2x and Figure S2B).

We localized the remaining escape mutations by diluting the 20 wells into two different media (+/-aTc) (Figure S2E&F). We concluded that 2/20 had mutated both TetR-Tup1 copies and 8 had mutated one but not both copies of TetR-Tup1. The remaining 10 grew only in the presence of aTc suggesting that those might have sustained mutations in LexA-hERLBD leading to constitutive repressive activity. This can happen due to mutations that yield a β-estradiol-independent-hERLBD, or truncations that leave the LexA moiety to repress transcription freely without β-estradiol. This suggested the third principle: truncations in the target must be selected against.

We placed a ribosomal skipping site and the *LYS2* gene after LexA-hERLBD, such that they would be transcribed as one mRNA but translated separately [22], [23] (Round 3). Since *LYS2* is required for growth in the absence of lysine, cells carrying early stop codons that create ligand-independent repressors would not survive. However, when grown in Synthetic Defined (SD) media lacking lysine, we observed increased growth rates at exactly the same stage along the growth curve (Figure 2C and Figure S2D). This suggested not stochastic mutations but a deterministic compensatory mechanism, suggesting a fourth principle: endogenous backup pathways that can substitute for the engineered growth control gene must be identified and removed.

In *S. cerevisiae*, expression of Pma2, the non-essential Pma1 homolog, can be increased under certain stress conditions[24] and we imagined that our acidic and nutrient-poor culture conditions might have caused compensatory expression of PMA2 (Figure 2B, Round 3). To test this idea, we constructed strains in which PMA2 was deleted, which prevented the adaptation we observed earlier (Figure 2C and Figure S2D). In this strain, only 12 of the 72 (16%) of wells showed fast growth within 5 days. Extrapolated to a 30-day campaign, this rate would correspond to roughly one-third of wells lost to escape if no beneficial target mutations arise in the interim, that is, a rate low enough to sustain evolution campaigns lasting 25 to 30 days. This was the timescale that our mathematical models (described in the next section) had predicted would be needed to reach improved target affinity.

Taken across the design iterations, each escape-suppression principle reduced the fraction of wells lost to escape by roughly an order of magnitude: from pervasive escape growth across the plate in Round 1 to 20 of 92 wells (22%) after adding the redundant Inverter, and to 12 of 72 wells (16%) in 5 days after introducing LYS2 truncation counter-selection and deleting the PMA2 bypass pathway. We note that these four principles are not specific to the Evolverator’s molecular components: any continuous evolution system that couples a non-essential protein function to growth through an engineered circuit will face the same classes of escape, and the same design principles should apply.

### Model-based analysis of system design to determine the optimal circuit architecture

In Evolverator, successful evolution requires coordination between processes that span molecular (affinity), cellular (growth) and population (evolutionary selection) scales. The right regimes for each scale are difficult to predict and time consuming to experimentally deduce. However, mathematical models can be used to quickly assess feasibility of potential Evolverator designs. We thus developed a mathematical modeling framework to guide circuit architecture decisions for this and future iterations of the Evolverator.

We used Ordinary Differential Equations (ODEs), a standard framework for modeling synthetic gene circuits, to describe expression levels and interactions of our components (Target, Mutagenesis, Inverter, Growth). However, ODEs cannot capture the probabilistic nature of evolution, such as the timing of random mutations and their subsequent effects on protein function. Since there were no other simple modeling frameworks that could interface with ODE-based models, we developed a mutagenesis model (Methods) and verified it using experimental data. This data consisted of fluctuation assays using *URA3* inactivation, in which we measured both mutation frequency in populations expressing different amounts of the mutator (Figure 3A, Figure S3A&B&C), and the time points at which these mutations arose during growth in liquid culture (Figure 3B). Our mutagenesis model combined ODEs to describe the expression level of the mutator with a poisson process to introduce mutations into *URA3*. Additionally, we added a growth model describing the growth dynamics of the two types of cells (wild type and inactivated *URA3*) in the culture. This allows us to correct growth and mutation rates based on the culture’s carrying capacity (the cell density at which the culture becomes saturated) as we are working with batch cultures. Model predictions accurately reproduced the experimentally observed distribution and dynamics of mutants (Figure 3A&B, Figure S3E&F).

**Figure 3.**
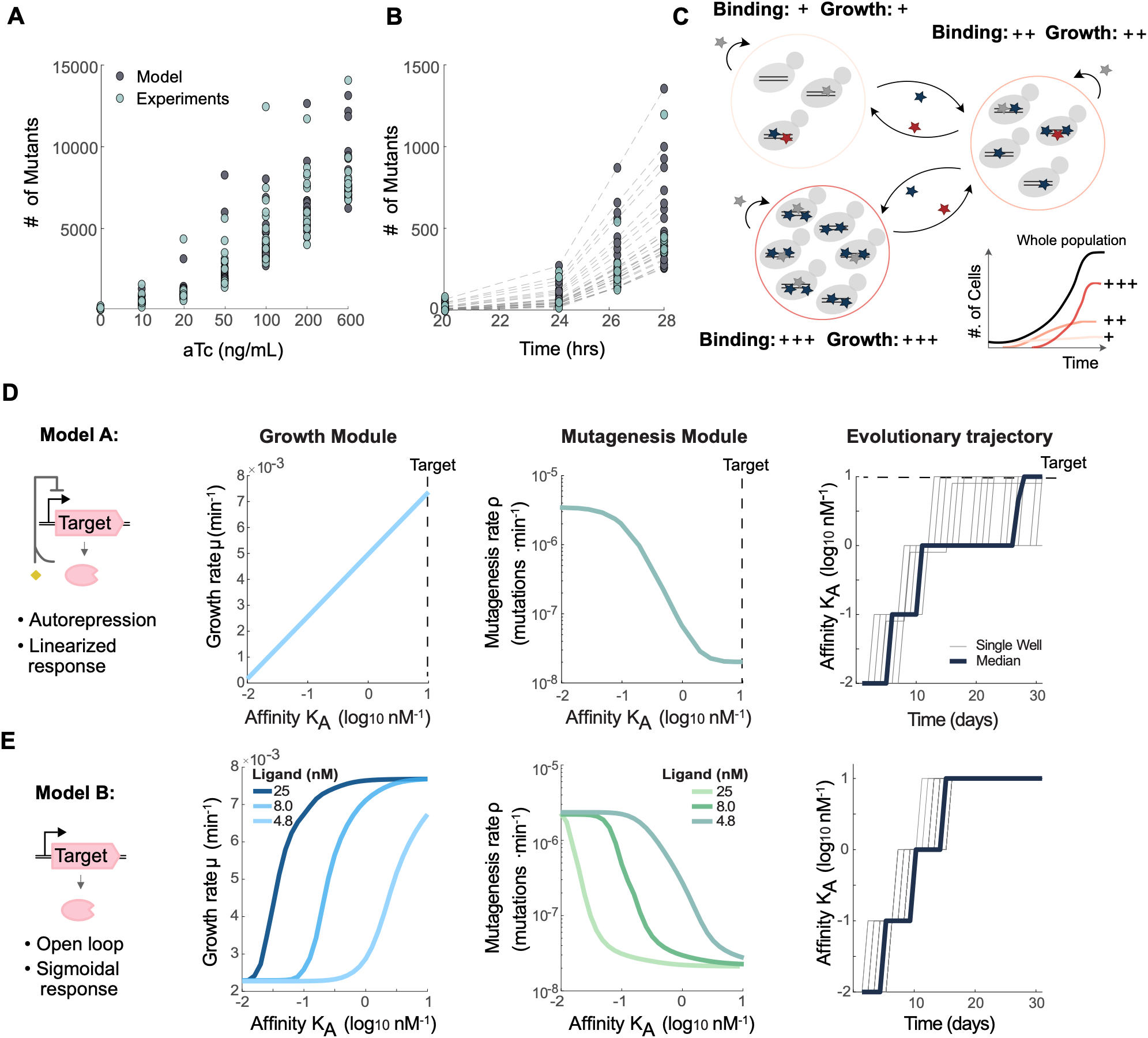
Model-based analysis shows the feasibility of the Evolverator design and suggests an Open Loop design with adjustable selective pressure for implementation. **A)** Experimental (green dots, individual experiments, n=12) and predicted (grey dots, individual simulations of the mutagenesis model, n=25) mutant cell numbers in 5’FOA fluctuation assays using a mutator expressed from an anhydrotetracycline (aTc) inducible promoter (see Methods). **B)** 5’FOA fluctuation assay with aTc = 200 ng/mL where the number of mutants was sampled over time from 6 independent wells; symbols as in (A), n=25 simulations. **C)** Simplified schematic of the model. Cells independently mutate their target receptor. Beneficial (deleterious) mutations increase (decrease) receptor affinity, creating discrete cell subpopulations over time with different growth rates. Neutral mutations are included but have no effect. Blue, red and grey stars represent beneficial, deleterious and neutral mutations, respectively. **D, E)** Predicted behavior of two Evolverator model versions (NF and OL), both simulated with their own optimized parameter sets for three design criteria: (i) precise control over the growth rate (left column) and (ii) the mutagenesis rate (middle), and (iii) the ability of the population to evolve the target receptor affinity (right). For iii: Grey lines: outcomes of single simulations, n=96; dark blue lines: dominant affinity, the most frequent affinity in the population. For E i-ii: different colors represent different ligand concentrations (see legend).

Then, to ensure that the mutagenesis component accurately represented hER rather than URA3, we extended the framework to account for a broader spectrum of mutations. We defined five classes containing both beneficial and detrimental effects and calibrated their probabilities using previously published mutagenesis data (Figure S3G and Methods). We integrated this probabilistic hER mutagenesis model with the ODEs describing other circuit components to create a complete model of the Evolverator system. Note that this model remains an abstraction of the full system: only the mutation probabilities for hERLBD are fixed, allowing us to evaluate whether the circuit design could achieve the intended behavior for any set of parameters, independent of specific molecular components.

We evaluated our circuit designs through simulations by examining (i) how changes in hERLBD ligand affinity influenced cell growth, (ii) whether cells shut-off mutagenesis after reaching the desired affinity and (iii) how many independent cultures reached our target affinity within a certain timeframe. Simulations begin with an isogenic culture. As each cell mutates its Target receptor, discrete groups of cells arise that differ by their affinity, and thus by their growth rate (Figure 3C). To mimic a 96-well plate, we ran 96 simulations. *In silico* cultures were diluted 1:250 at saturation, and each dilution-to-saturation cycle was considered one growth period. Because of these dilution steps as well as the carrying capacity, time and ‘population state’ at which a beneficial mutation occurs influences whether this cell can outgrow the other cells. For example, beneficial mutations arising in a dense culture are unlikely to have sufficient time for outgrowth.

We first tested a design in which the Target also repressed its own expression, since we and others had shown that autorepression can linearize dose responses[14], [25]. This could create a linear affinity-growth relationship, enabling evolution without any manual interventions to adjust selective pressure. Simulation of this model for an optimized set of parameters indeed produced a linear relationship between affinity and growth as well as the desired inverse relationship between mutagenesis and growth (Figure 3D). However, evolution slowed down as soon as the mutation rate decreased, since cells with one beneficial mutation took longer to generate additional mutations needed to increase affinity.

On the other hand, simulations with constitutive Target expression resulted in a sigmoidal affinity-growth rate relationship, which allowed us to change the required affinity for a given growth rate by changing the ligand concentration (Figure 3E). Doing so accelerated evolution by keeping the mutation rate high and giving subsequent higher affinity mutations a growth advantage. Thus, our model identified the sigmoidal regime’s advantage over the linear autorepression alternative before either architecture was built in cells.

These simulations showed that we could expect to obtain improved Target variants every 5-7 days, suggesting that three-week evolution campaigns, comparable to the time frame for previous automated, continuous evolution experiments [26], might produce improved variants. In addition, we also found that our model was consistent with the order of escape mutations we found: small changes in the parameters belonging to the inverter module, i.e., TetR-Tup1, restored full growth, but the LexA-hER module was less sensitive to (parameter) mutations (Figure S3J).

### Evolution of receptors with increased affinity for different estrogen derivatives

We used the escape-proofed Evolverator_hER_ strain (Figure 4A) to carry out separate evolution campaigns lasting 28, 22 and 21 days to increase affinity of hERLBD towards three estrogen derivatives (β-estradiol, estrone and estriol). Cells were grown in 96 well plates and diluted 1:250 whenever most wells reached saturation (each dilution → saturation is one growth period). The cells also contained a fluorescent protein (Citrine) expressed from a LexA-hERLBD repressible-promoter. If hormone binding affinity of hERLBD increased, we would expect to see reduced Citrine expression. As a positive control, we set up wells grown in presence of high aTc, allowing full Pma1 expression. These wells had no restriction on their growth and showed no repression of Citrine due to lack of hormone ligand. This allowed us to monitor the state of evolution by comparing the growth rate and the fluorescence observed in each well to the control wells. We could then adjust selective pressure during the campaign by reducing concentration of the hormone (Figure S4A).

**Figure 4.**
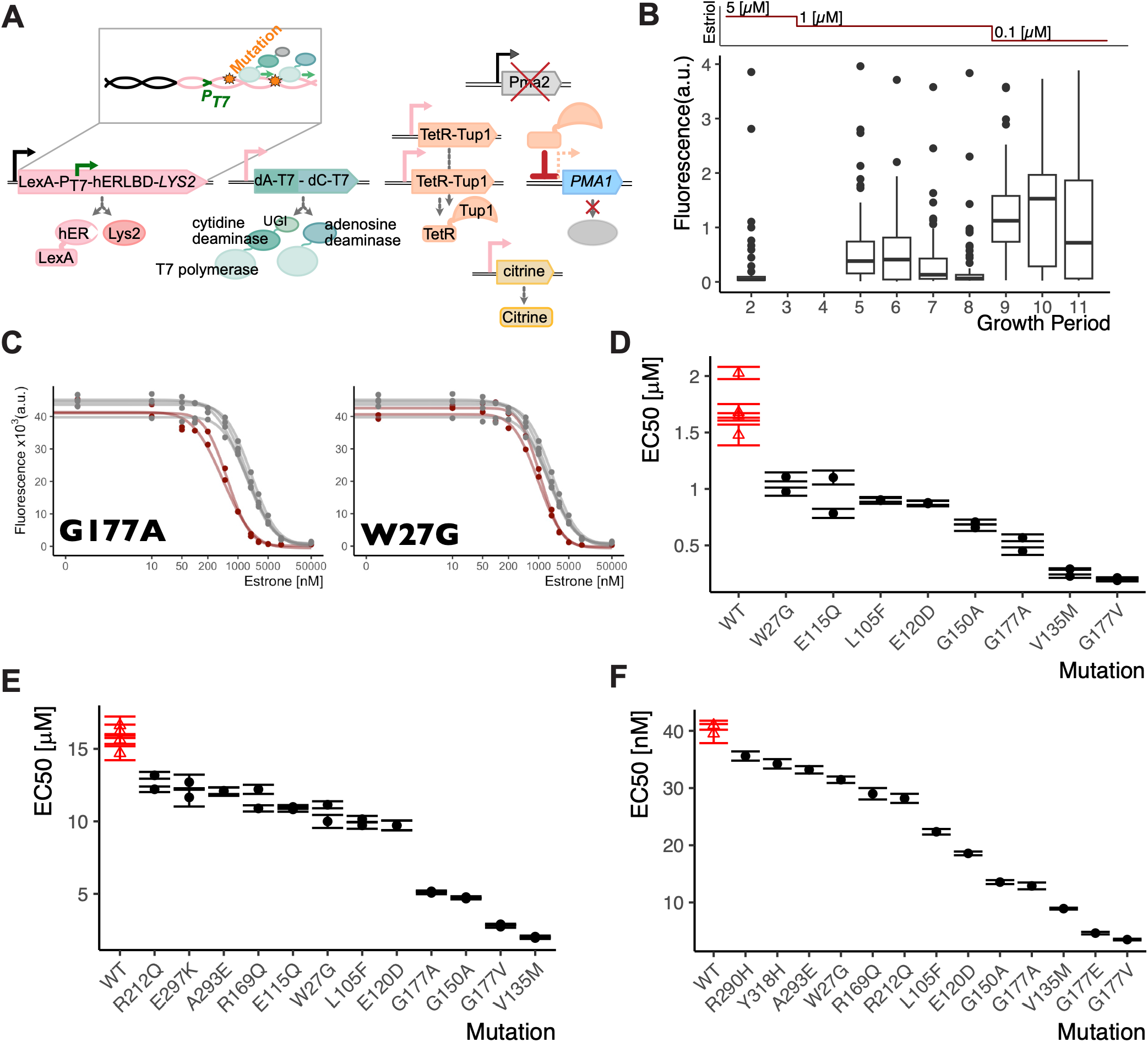
The Evolverator_hER_ identifies mutations that increase ligand affinity of hERLBD towards hormones. **A)** Final molecular implementation of the Evolverator_hER_ (Y4037). Active (solid) and inactive (dashed) promoters and the proteins produced are shown. Promoters are the same color as the proteins that can repress them. Proteins that are not produced are depicted in grey. LexA-hERLBD acts as a repressor when bound to hormone (e.g. β-estradiol). The mutators initiate transcription at the T7 promoter placed between the coding regions of LexA and hERLBD. Deaminases introduce mutations (orange stars) as the T7 polymerase transcribes. The mRNA produced by T7 polymerase is not translated and is degraded by the cell. **B)** State of evolution throughout the estriol campaign. Citrine fluorescence per well was collected with flow cytometry and normalized to the median of one positive control well without hormone. Values below 1 indicate increased repression by LexA-hERLBD. Lines: population medians; boxes: 25^th^ to 75^th^ quantiles. Estriol concentrations during growth periods 1-3: 5µM, 4-8: 1µM, 9-11: 0.1µM. **C)** Examples of the two mutations identified in the campaigns. Strains bearing P_7SulA.1_ expressed Citrine and mutated (red) or original (grey) LexA-hERLBD were grown in YPD and varying estrone concentrations. Dots: population medians of Citrine fluorescence measured by flow cytometry; error bars: 25^th^ to 75^th^ quantiles; lines: fitted dose response curves (see Methods). **D-F)** Comparison of EC_50_ values for estrone (D), estriol (E), and β-estradiol (F) of mutant LexA-hERLBD versions tested as in (C). Fitted dose response curves were used to calculate EC_50_ values. Error bars: estimated standard error of the estimated parameter values.

During these campaigns, at constant hormone concentrations, median population fluorescence decreased over time, indicating increased repression by mutant LexA-hERLBD versions, and fluorescence increased again when we reduced hormone concentration to increase selective pressure (Figure 4B and Figure S4B,C). We terminated each campaign once almost all evolution wells had the same growth rate as positive control wells and many evolution wells showed low fluorescence. For the estrone and estriol campaigns, this was the end of the 8^th^ growth period (88/88 and 87/88 wells with 1µM hormone showed fluorescence below control wells; Figure S4D). For the β-estradiol campaign, we collected samples from the 11^th^ growth period but noticed that many wells had evolved constitutive, hormone-independent repression (reduced fluorescence observed in the well with and without β-estradiol). We therefore replaced these wells with cells from the 6^th^ growth period for sequencing (Figure S4E).

Sequencing revealed 28 (estrone), 21 (estriol), and 53 (β-estradiol) unique single mutations in the hERLBD that presumably increased the affinity of the hERLBD to those ligands or otherwise increased repression by LexA-hERLBD (Supplementary Tables 1A-C). We cloned 20 randomly chosen unique single mutations into LexA-hERLBD and tested the repression efficiency and ligand affinity with dose response experiments (Figure 4C and Figure S5). We fitted a simple four-parameter model to the dose response data and estimated EC50 values (Figure 4D-F and Methods). For estrone, estriol, and β-estradiol, 8/17, 12/17 and 13/18 mutations tested decreased EC50 values with a maximum 11.5 (0.1)-fold improvement in EC50 (Supplementary Table 1D).

None of the validated clones in the three campaigns match the ligand-contacting residues targeted in prior directed evolution efforts on the hERLBD [27], [28]. Instead, identified mutations mapped to positions that modulate receptor function via allosteric, scaffolding, and dimerization mechanisms (Figure S5D). We assign structural positions using full-length estrogen receptor (ESR1) (our construct began at residue 266 in the hinge region). Our mutations fall into two structural classes.

One class sits near the binding pocket and enhances ligand response. Position 177 (ESR1 G442, helix H8) was the most frequently mutated; four different substitutions (G177A, G177V, G177E, G177R) were recovered across campaigns, with G177V showing 11.5-fold decreased EC50 for β-estradiol, likely by rigidifying the H7–H10 helical core. V135M (V400M, H5–H6 loop), the strongest single mutant outside position 177 (4.6-fold decreased EC50 for β-estradiol), lies adjacent to R394 and E353, which together with a conserved water molecule hydrogen-bond the estradiol A-ring hydroxyl[29]. G150A (G415A, helix H7) constrains the geometry of the binding pocket with a 3.0-fold EC50 decrease. Residues 135 and 177 were previously linked to hormone-dependent dimerization and ligand-dependent activity[30] and indeed mutations improved EC50 comparably for all three hormones. The second class (W27G/ W292G in the hinge; R169Q/R434Q on helix H8; and R212Q/R477Q at the H10/H11 dimer interface) modulates receptor conformation or dimerization geometry, with more modest and hormone-specific EC50 effects. Together, these results demonstrate numerous evolutionarily accessible mutational paths to improved receptor function beyond the canonical ligand-contacting residues identified by more classical structure-guided medicinal chemistry or saturation mutagenesis of the ligand binding pocket.

We noticed that while certain mutations came to dominate wells, multiple low-frequency mutations could also be found in each well. For example, the three campaigns also yielded double and triple mutations, albeit at low frequencies (Supplementary Table 1E). We tested nine double mutations to see their effects (Figure S6A) on LexA-hERLBD. For β-estradiol, two double mutants (G177V, E324K and G177A, D109N) showed increased affinity, with the latter showing a synergistic increase over the single mutation controls (Figure S6A). Out of the remaining seven, one did not affect hormone affinity and six reduced or completely abolished it. Additionally, at the end of the β-estradiol campaign, each well had an average of 4.7 unique single mutations and 39 out of 88 wells had at least two unique multiple mutations (Figure S6B,C). In other words, neutral and deleterious mutations seem to persist in evolving populations at low frequency, demonstrating the potential of Evolverator to find solutions that require multiple beneficial mutations. Solutions with double or more mutations were probably not observed in the hERLBD campaigns (except for G177A, D109N) likely due to the abundance of easily accessible single mutations (at least three separate mutations of G177) that substantially improve affinity.

### Evolution of lac repressors with enhanced repression

We next decided to evolve a different protein binding interaction and moved to our second Target, LacI. LacI is a prokaryotic DNA binding protein, and its binding can be controlled by IPTG. However, our goal was not to evolve LacI-IPTG binding, but rather LacI-DNA binding to make LacI a better transcriptional repressor of a yeast promoter (i.e. reduce leaky expression)[15]. The Evolverator_LacI_ cells, Y3869, carried modifications to the design (Figure 5A). Citrine-P_T7_-LacI was used as the Target, allowing P_T7_ and the yeast promoter to be well-separated, preventing any potential transcriptional interference. Because truncations within LacI would likely not yield better repression, we did not include a *LYS2* gene. Weak LacI repression (Figure S7A) allowed TetR-Tup1 expression, such that only little *PMA1* is expressed, and cells proliferated slowly. The Mutator subsystem was placed under the control of LexA-hERLBD, and we used β-estradiol to suppress mutator expression whenever the strain was handled outside of evolution campaigns. A fluorescent protein (mApple) was placed under the control of a LacI-repressible promoter to track repression efficiency during evolution.

**Figure 5.**
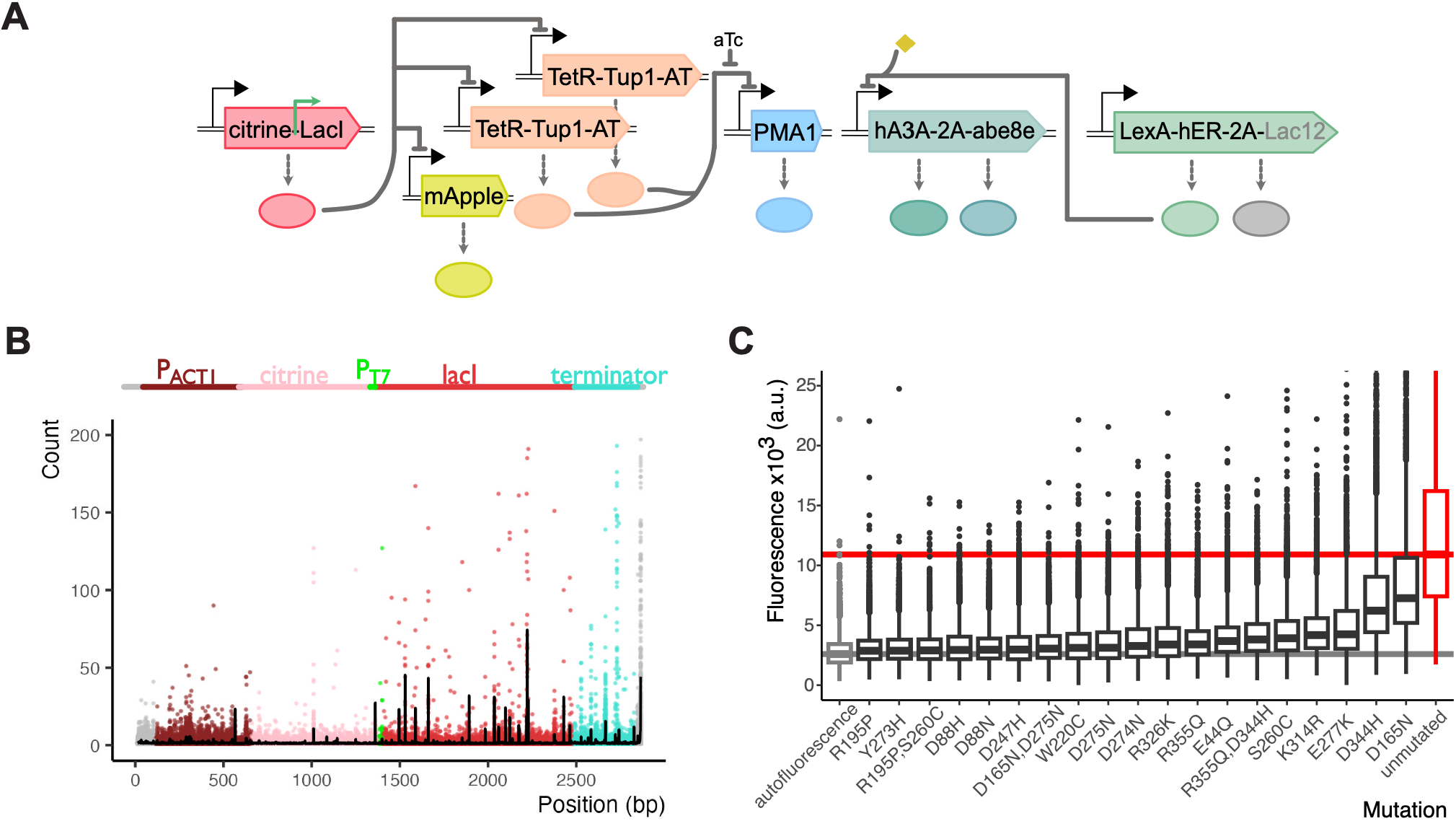
Evolving increased transcriptional repression by LacI. **A)** Molecular implementation of the Evolverator_LacI_ (Y3869). Symbols as in Figure 4A. **B)** Positions of DNA mutations at the end of the evolution experiment with strain Y3869 (see Methods). Dots: mutation counts per well (n=92) at each base pair position (1 is the first sequenced base); black lines: mean number of mutations at each position. **C)** Repression efficiency of LacI versions found in (B). Mutations were introduced into wild type LacI separately and strains expressing the mutants and LacI repressible Citrine were grown in rich medium (Y3965,3968:3972,3974,3976,3977,3982:3991). Lines: median of the population; boxes: 25^th^ to 75^th^ quantiles; autofluorescence: parent strain without Citrine (Y3316); unrepressed: strain without any LacI (Y3542); unmutated: original LacI sequence.

We used Evolverator_LacI_ to carry out two parallel evolution campaigns. Plate 1 had only rich medium. Plate 2 had a small amount of aTc (4 ng/mL) until the end of the 5^th^ growth period; after which we removed aTc to decrease *PMA1* expression and thus increase selective pressure. Both plates included positive control wells where cells grew at full speed in presence of high aTc. We sampled the wells every growth period for flow cytometry measurements of mApple. We normalized the median mApple fluorescence in each well to the average median of the control wells (Figure S7B), such that a value lower than 1 indicates increased repression strength of LacI in a well. We stopped evolution after the 8^th^ (plate 1) and 10^th^ (plate 2) growth periods when almost all evolution wells grew as fast as the control wells and mApple fluorescence was below 70% of control in 30 (plate 1) and 31 (plate 2) wells.

With sequencing we detected mutations in several regions of the Target, most frequently within the LacI sequence (Figure 5B). In LacI we identified 25 unique nonsynonymous mutations (Supplementary Table 1F). In 14 wells, two mutations occurred simultaneously, giving us a total of 5 unique double mutations (Supplementary Table 1G).

We incorporated three sets of double mutations and 16 randomly chosen single mutations into wild type LacI and tested them by quantifying repression of a LacI-repressible promoter expressing Citrine. All 19 mutations increased LacI repression compared to the original, unmutated LacI, and there were no false positives (Figure 5C). Lack of any false positives suggests that all 25 single mutations found in the evolution campaign probably increased LacI repressor activity. For the double mutations, repressed Citrine levels were similar to those of the stronger single mutations, and we did not detect synergistic effects.

20 of the recovered LacI mutations lie in the repressor’s core domain, which contains the inducer-binding pocket and the dimerization interface and two lie in the DNA-binding domain (D8N and E44Q) and two in the tetramerization domain (D344H and R355Q) (Figure S7E). Four core-domain mutations (D247H, D275N, W220C, D274N) abolished IPTG response entirely (Figure S7C&D), suggesting they shift the allosteric equilibrium so strongly toward the DNA-bound conformation that IPTG can no longer release the repressor [31]. Nine mutations preserved normal IPTG response, enhancing repression without disrupting the allosteric switch, and two (D88N, D88H) showed weakly responsive intermediate behavior. This variation in IPTG-response is expected, since during evolution with did not create any selective pressure to maintain IPTG response. This distribution defines an evolutionarily accessible dataset on allosteric decoupling: some mutations lock the repressor onto DNA, while others altered binding affinity while preserving inducibility.

Comprehensive mutational studies were already performed on LacI in *E*.*coli* [32], [33], [34], [35], [36], [37], [38], [39], including a recent saturation mutagenesis study[40], which provides a comprehensive baseline against which the Evolverator’s mutations can be compared (see Supplementary Table 1H for details). Five mutations showed unchanged, diminished or abolished function in the saturation mutagenesis dataset in *E*.*coli* (D247H, R195P, D344H, D88N, D88H) yet repressed more strongly in yeast. Although repression by prokaryotic repressors at synthetic yeast promoters has been exploited for decades, that repression differs from *E. coli* repression in ways that are still not fully mechanistically characterized, making it plausible that substitutions that attenuate repression in the bacterial host can enhance it in the yeast context.

### Validating the *in silico* evolution models to help design future evolution campaigns

A validated mathematical model can play an important role in the design of future Evolverator strains. We therefore parametrized our Evolverator Model with data from independent experiments on the individual Evolverator modules. Our estimated parameters captured the joint experimental data well (Figure 6A,B and Figure S8, S9), which we used to create Evolverator Models specific for hER and LacI (Methods). To evaluate their performance, we used them to simulate evolution experiments with the same sequence of dilutions used in the experimental campaigns. To compare *in-silico* and experimental results, we compared mid-points: time points at which individual wells reach half-maximal cell density per growth period. Simulations captured the experimental mid-points (Figure 6C), both in terms of the spread between the wells and of median. An exception is the initial mismatch for estrone binding evolution. Experimentally, all evolving wells grew comparably to positive controls for the first three growth periods, and our model recapitulated this behavior only if our simulations posited that all wells developed a mutation (e.g., one that increased LexA’s repression efficiency or stability) very early, during initial diversification or at the start of the first growth period (Figure 6C, S10E). Given the rather simple modeling framework – it neither represented cell-to-cell variation nor attempted to capture escape mutations – the realism of predicted dynamics and outcomes of evolution was encouraging.

To identify critical factors for future Evolverator designs, we simulated how sensitive *in silico* evolution campaigns would be to changes in the hER Evolverator Model parameters (Methods and Figure S11A). We saw that changing protein production rates or repression coefficients – while keeping the other parameters fixed – easily lead to a non-functional system (Figure 6D). This might be because in such an interconnected, complex system, the ratios between parameters, rather than their absolute value, is important for a system’s behavior. To identify all such correlations between our parameters we sampled the parameter space using Markov chain Monte-Carlo (MCMC) and calculated parameter correlations (Methods). There were several strong positive correlations (Figure 6E, Figure S11B) suggesting that concentrations of certain species need to match precisely to obtain correct input-output relationships between Evolverator modules. This suggests that future Evolverator designs should ideally use system components that can each be tuned with exogenous chemicals (like TetR-Tup1 could be with aTc). This makes it easier to adjust parameters (like expression level or repression strength) that need to match precisely and optimize the system behavior, compared to designs that require genetic modifications to adjust these parameters.

Our parametrized models can be used to transfer the Evolverator concept to other evolution scenarios and binding interactions. For example, simulations using the LacI-parametrized model, but starting from a weaker initial repression efficiency, suggest that a general strategy to evolve stronger transcriptional repressors is to express them under a stronger promoter than that used for LacI and to progressively reduce the fraction of the repressor in its active, DNA-binding form (Figure 6F). This can be achieved either by adding a small molecule that interferes with DNA binding or by using an inducible promoter to modulate expression.

We can also use our modeling framework to determine *in silico* the requirements (e.g. promoter strength, reaction rates) for new binding interactions that we want to incorporate in our system. For example, Figure 6G shows a successful simulated design in which stronger protein-protein interactions can be evolved by fusing both proteins to two parts of a two-hybrid system, such that the system functions only when they are bound together [41]. In this setup, we cannot compensate for initial weak protein–protein interactions by simply increasing the expression of one partner, as we could with small molecule ligands (due to cellular burden of overexpressing proteins). Instead, we achieved succesful *in silico* evolution by increasing the repression efficiency of the Target complex to be higher than that of our original LexA-hER target. This example represents how we can use the model to check whether a new binding interaction can directly function within the Evolverator or whether the system components require modification. The ability of the model to specify required promoter strengths, reaction rates, and module stoichiometries for a new target before any strain is built, is what lets it prescribe, rather than merely evaluate, the circuit modifications needed to retarget the platform in the future.

## DISCUSSION

We developed *in vivo* continuous directed evolution for non-essential eukaryotic proteins in yeast, which operates in high throughput liquid cultures without special equipment. Evolverator cells continuously mutagenize and screen variants of a target protein while a rationaly designed genetic circuit drives faster proliferation in cells with variants approaching the desired function. We used Evolverator cells to enhance two protein binding interactions: a nuclear hormone receptor (human estrogen receptor), to increase interaction with three small-molecule hormones, and a repressor (LacI), to increase repression of an engineered eukaryotic promoter.

Most mutations identified in hERLBD and LacI likely enhance stability of the active protein conformation rather than directly altering ligand binding or intrinsic affinity. This bias may reflect the relatively high starting affinity of both proteins, making further improvements more accessible through conformational stabilization. Like any other reporter-based evolution system, Evolverator selects for apparent functional output rather than specific molecular properties such as true binding affinity, a limitation that should be considered when designing future Evolverator circuits.

The four design principles we identified for escape-resistant evolution (separation of growth control from metabolic burden through asymetrically inherited proteins, genetic redundancy in key components, selection against mutations that truncate the evolutionary target, and removal of endogenous bypass pathways) are not specific to the Evolverator’s molecular implementation. They address failure modes inherent to any system that couples a non-essential protein function to growth through an engineered circuit, and we expect them to apply broadly.

Moreover, we used a model-> design-> validation approach to improve our gene circuit design and gain a better understanding of the system. The close match between our model simulations and growth dynamics observed in our evolution campaigns validates our model and helps us to predict adjustments needed for new protein targets and classes. This makes the system more readily adaptable. In the future, the models we developed here can be expanded to include escape mutations to optimize experimental protocols to limit their occurrence or aid in the design of selection/counter-selection schemes. For example, in the LexA-hERLBD campaign, counter selection (selecting for growth in the absence of the hormone ligand) could be used to “prune” mutants with constitutively active repression.

A practical question for any evolution platform is how much effort is required to adapt it to a new target. For the Evolverator, the concrete steps are: (1) engineer a construct that couples the target protein’s function to transcriptional repression of the Inverter module; (2) use the parametrized mathematical model to compare the target’s expected affinity range and expression levels against the circuit’s functional parameter space; (3) where the expected parameters fall outside that space, use the model to identify which circuit modifications (e.g. promoter strength, copy number, binding-site count, module stoichiometry, degron choice) and move them inside, and iterate steps (2) and (3) *in silico* until the design is predicted to work; and (4) build the strain and verify module behavior using the informative module approach described in here (Methods). For a molecular biologist experienced with yeast genetics, we estimate this process would require two to three months from design to a functional evolution strain, with the modeling steps taking days rather than weeks. This compares favorably to OrthoRep, which requires designing a growth-coupled selection and cloning the target onto the p1 plasmid. This is simple when the target is essential for growth, but without a gene-circuit approach like Evolverator, it is not possible to establish for non-essential proteins. The Evolverator’s modeling framework thus offers a distinct advantage: the model does not merely evaluate whether a new target is compatible with the circuit, but it prescribes the circuit modifications required to make it compatible before strain construction begins.

As we were preparing this manuscript, a similar continuous evolution system named Optovolution was published [11]. Optovolution couples a light-regulated transcriptional regulator to expression of the S-phase cyclin Clb5 and evolves it for improved light sensitivity, reduced leakiness, altered spectral response, as well as a two-input logic gate. These are impressive demonstrations of selecting for optogenetically defined protein behaviors, which the Evolverator did not address but could easily accommodate. Most significantly, the two systems differ in how they establish the phenotype-fitness link. Clb5 gates entry to S-phase and thus blocks DNA replication. This will inevitably disrupt cell physiology and lead to off-target escape mutations. Indeed, campaigns were conducted on solid agar (limiting throughput and automation); a brief liquid-culture trial was reported but not pursued further, consistent with escape mutations being harder to contain in continuous liquid growth. Additionally, DNA replication is required for introducing mutations into the target gene with the error-prone polymerase employed in Optovolution. Therefore, the growth control and mutagenesis components are inherently at odds with each other in this system. On the other hand, the Evolverator’s Pma1-based growth control decouples fitness from the cell cycle and mutagenesis and operates in standard liquid culture over weeks. The two systems are therefore different in scope: Optovolution demonstrates light-directed, low throughput evolution on solid medium; the Evolverator demonstrates transferable escape-suppression principles and predictive modeling for sustained evolution of binding-driven functions in high-throughput liquid culture.

Evolverator is complementary to AI-guided protein engineering rather than an alternative to it. Computational methods propose starting libraries and prioritize positions for diversification; the Evolverator returns quantitative variant-fitness data that is generated in a eukaryotic cellular context. This kind of data is crucial for building accurate prediction capabilities on new targets, which purely computational approaches cannot produce on their own. It is important to note that in our evolution campaigns we did not start from diversified libraries and yet still identified many successful mutants. Starting from computationally designed, focused libraries will likely improve outcomes of Evolverator campaigns significantly in the future.

Overall, Evolverator’s efficiency in screening mutants and ability to link different functions to a stringent and tunable growth selection, as well as the faithfulness of the accompanying mathematical models, mean that its application could be extended to other target classes such as enzymes, G-protein-coupled receptors and even ion channels. In a host like *S. cerevisiae* that can express many eukaryotic proteins, the ability to rapidly engineer systems that can evolve non-essential proteins has the potential to generate custom proteins with wide applications in biotechnology and biomedicine.

## Author Contributions

AA performed the wet lab experiments, EB the computational studies, and contributed equally to this work. RB described the original design and advised on the work. AA, EB and RB wrote the manuscript, and all guarantee the integrity of the work.

## Acknowledgments

We are very grateful to Joerg Stelling for contributions to conceptualization of the work, supervision, funding acquisition, and feedback and work on the manuscript. We thank Erica Geneletti for preliminary computational work, and Claude Magnard Lormeau, Alejandro Colman-Lerner, Harmit S. Malik, Alexander Mendenhall, and Andrew Ellington for useful discussions.

## Methods

### Lead Contact

Further information and requests for resources and reagents should be directed to and will be fulfilled by the lead contact Asli Azizoglu.

### Materials availability

All plasmids and strains generated in this study are available from the authors upon request.

### Data and code availability

All original code, all experimental data including raw flow cytometry measurements, colony counts and culture density measurements are deposited at the ETH Research Collection (https://doi.org/10.3929/ethz-c-000803791) and are publicly available. All next-generation sequencing data is available publicly through the SRA under BioProject Accession number PRJNA1503740.

Any additional information required to reanalyze the data reported in this paper is available from the lead contact upon request.

### Experimental model and study participant details

All *S. cerevisiae* strains used in this study can be found in Table S1I. All strains used were based on a haploid derivative of the BY4743 strain (MATa *his3Δ leu2Δ met15Δ ura3Δ lys2Δ*). Cells were grown at 30°C with shaking (275rpm) and stocks were kept in 15% glycerol at −80°C.

*PMA2* open reading frame (ORF) deletion was performed using the CRISPR-Cas9 protocol previously described[42]. Briefly, the guide RNA was designed using the sequence AATAACCGTCGAAGGGGGAA for the variable region, was produced by Thermofisher, UK, and transformed together with a plasmid carrying inducible Cas9. 10µM β-estradiol in liquid culture was used to induce the expression of Cas9 and create a double stranded break. The sequence GACAGACGCTGCTCAAATAACCGTCGAAGGGGGAATAATTTTAAACATCATCCATCTGATTTTTTTTCTT was used as the repair template to remove the region between 296bp before the start codon and the end of the ORF (including the stop codon).

When constructing the strain where P7tet.1 replaces the endogenous promoter of PMA1, the same CRISPR-Cas9 based protocol was used[42]. The guide RNA was designed using the sequence GTTAATAATTAATTAATTGG for the variable region. A PCR fragment containing P7tet.1 and an antibiotic marker (Nourseothricin (Werner BioAgents, clonNAT)) was generated from plasmid P2375 with the oligos atgctgaagaggatgatgaagaggatgatgtatcagtcattttcccttttccctt and GATAAATTTTTTCTTTAACAATCGTTAATAATTAATTAATagcttgccttgtccccgcc. This was transformed for homologous recombination directed replacement of the endogenous promoter during the second transformation.

### Plasmids

All plasmids used in this study are detailed in Table S1J and relevant sequences are given directly in Table S1K. The DH5 α *E*.*coli* strain grown at 37°C was used for production of all plasmids. All relevant sequences can be found in Table A12. All plasmids with auxotrophic markers were constructed using the pRG backbones with isothermal assembly[43]. Inserts were generated by PCR on existing plasmids using oligos synthesized by Microsynth, Switzerland or Thermofisher, UK. The pRG backbone plasmids integrate into the genome as a single copy construct. They were linearized using the restriction enzyme AscI before transformation. In case of the LexA-PT7-hERLBD-2A-*LYS2* plasmids, the backbone contained no auxotrophic marker and selection was based on the *LYS2* present within the open reading frame. Plasmids containing antibiotic resistances were based on the pFA6 backbone[44].

The cytidine deaminase hA3A-T7P fusion used for mutagenesis in various strains had a Uracil DNA glycosylase inhibitor (UGI) peptide fused to its C-terminus, since this was shown to increase mutation frequencies by preventing cellular repair machinery from excising the uridine generated from cytidine deamination[20], [45].

In plasmids where both mutator fusions hA3A-T7P and abe8e-T7P were present, they were separated by the intra-ribosomal self-cleaving peptide sequence F2A. This sequence allows two independent proteins to be translated from a single transcript[46].

### Chemicals and Media

YPD was prepared with 1% yeast extract (Thermofisher, 212720), 2% bacto-peptone (Thermfisher, 211820) and 2% glucose (Sigma, G8270). Synthetic (S) media without lysine contained 0.17% yeast nitrogen base (without amino acids and ammonium sulfate) (BD Difco, 233520) with 0.5% ammonium sulfate (Sigma, 31119) as nitrogen source, a complete complement of amino acids except for lysine, and the supplements adenine and uracil. SD plates with 5’FOA contained all supplements listed for SD media above, with the addition of lysine and 1.5mg/mL 5’FOA.

aTc was purchased from Cayman Chemicals (10009542) and prepared as a 4628.8ng/mL (10mM) stock in ethanol for long term storage at −20°C and diluted in appropriate media for experiments as necessary. All hormones and hormone substitutes (β-estradiol (CAS number 50-28-2), Estrone (CAS number 53-16-7), Bisphenol A (CAS number 80-05-07), Tamoxifen (CAS number 105-40-29-1), Estriol (CAS number 50-27-1), and Genistein (CAS number 446-72-0)) were acquired from Sigma and dissolved in 70% ethanol for long term storage at −20 °C. They were further diluted in appropriate media for experiments as necessary.

### Dose response experiments

Cells were grown overnight in appropriate media and diluted 1:200 into experiment conditions with varying inducer concentrations in 96 well plates with 250µL media. Plates were shaken at 210rpm overnight at 30°C. Cells were diluted 1:20 into the same conditions in the morning, and fluorescence was measured using flow cytometry after 7 hours.

### Flow cytometry

Samples were diluted in PBS and measured using an LSRFortessa LSRII equipped with a high-throughput sampler. PMT voltages for the forward and side scatter measurements were set up such that the height of the signal was not saturated. Citrine fluorescence was quantified using a 488 nm excitation laser and a 530/30 nm emission filter. mApple fluorescence was quantified using a 561 nm excitation laser and a 610/20 nm emission filter. PMT voltage for these channels were set up such that the signal from a control strain expressing high levels of Citrine or mApple did not saturate the measurement device, except for basal level measurements in Figure 5C, where PMT voltage for the Citrine channel was increased to maximum. Side scatter was measured using the 488 nm excitation laser and 488/10 nm emission filter.

### Fluctuation Assays

Cells were cultured overnight without uracil and diluted into a 96 well plate with rich media with 600ng/mL aTc at a concentration of 1000 cells/well. They were shaken for 30 h in continuously growing cultures at 30°C. Cultures were then diluted in PBS and plated on SD 5’FOA plates. The dilution rate was decided based on preliminary experiments, such that single colonies could be observed when 100µL of the diluted culture was plated as a spot and allowed to dry. Where more mutations (and therefore more colonies) were expected, the dilution rate was higher. At least 10 parallel cultures of each strain were plated. Based on the number of colonies that grew on this selective plate, we calculated mutation frequencies and confidence intervals using a Newton-Raphson type algorithm implemented in the R package rsalvador. We used the “newton.LD.plating” function, which takes into account the dilution step before plating. The confidence intervals were calculated using the “confint.LD.plating”, again taking into account the dilution step.

### Liquid Growth Assays

Cells were grown overnight at 30°C in the same liquid culture the experiment was to be performed in. They were diluted into 96-well plates with transparent bottoms (Enzyscreen, CR1496dg) with various inducer concentrations where appropriate. The total liquid volume in each well was 250µL. Plates were covered and shaken (250rpm) at 30°C by the Growth Profiler 360. Pictures of the bottom of the plate were taken every 20 or 30 minutes depending on the experimental setup. Growth Profiler Viewer 360 software was used to automatically quantify culture density in each well from these pictures. Culture density values correspond to this calculated value minus the lowest, background value recorded in a given well. This correction sets the lowest recorded value as 0 for ease of plotting.

### Next Generation Sequencing

At the end of each evolution experiment, 200 µL of liquid culture from each well was transferred to a 96-well filter plate with 0.2µM PVDF filters (Sigma Aldrich, CLS3508) and centrifuged for 1 min at 1500g. The remnants in the filter were resuspended in 150µL of 0.2M LiAc+1%SDS solution and incubated at 70°C for 30 minutes. 150µL pure ethanol was added and the plate was centrifuged again for 1 min at 1500g. The remnants were resuspended in 60µL double distilled, sterile water. This gDNA extraction protocol was adapted from[47].

1 µL of the gDNA solution was added to 50µL of NEBNext Ultra II Q5 master mix for PCR. The forward primers had unique barcodes for each well of the experiment and the reverse primers contained unique barcodes for each evolution experiment. The PCR product was run on 1% agarose gel using the SYBR Safe DNA gel stain (Thermofisher, S33102) and gel purified using a gel extraction kit (Qiagen, 28706×4). Samples were submitted for SMRTBell library preparation and PacBio sequencing by the Genomics Facility Basel.

### Data analysis

Dose response curves were generated using a 5-parameter log-logistic function as explained previously[14].

The raw NGS reads were demultiplexed based on the unique well and experiment barcodes using the axe-demultiplexer and the axe-demux command. The demultiplexed reads were aligned to the reference sequence using the bowtie2 command with default settings within the bowtie2 package. Downstream analysis was performed using a custom R script.

Slopes of growth curves were calculated as 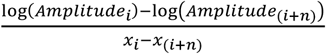 where *xi* is the time at which a measurement was taken. Depending on whether measurements were taken every 20 or 30 minutes, n is either 16 or 24 such that the two measurements are separated by 480 minutes. Maximum slope recorded over the growth curve is reported as the maximum growth rate (MaxGR).

### Overview of the modeling processes

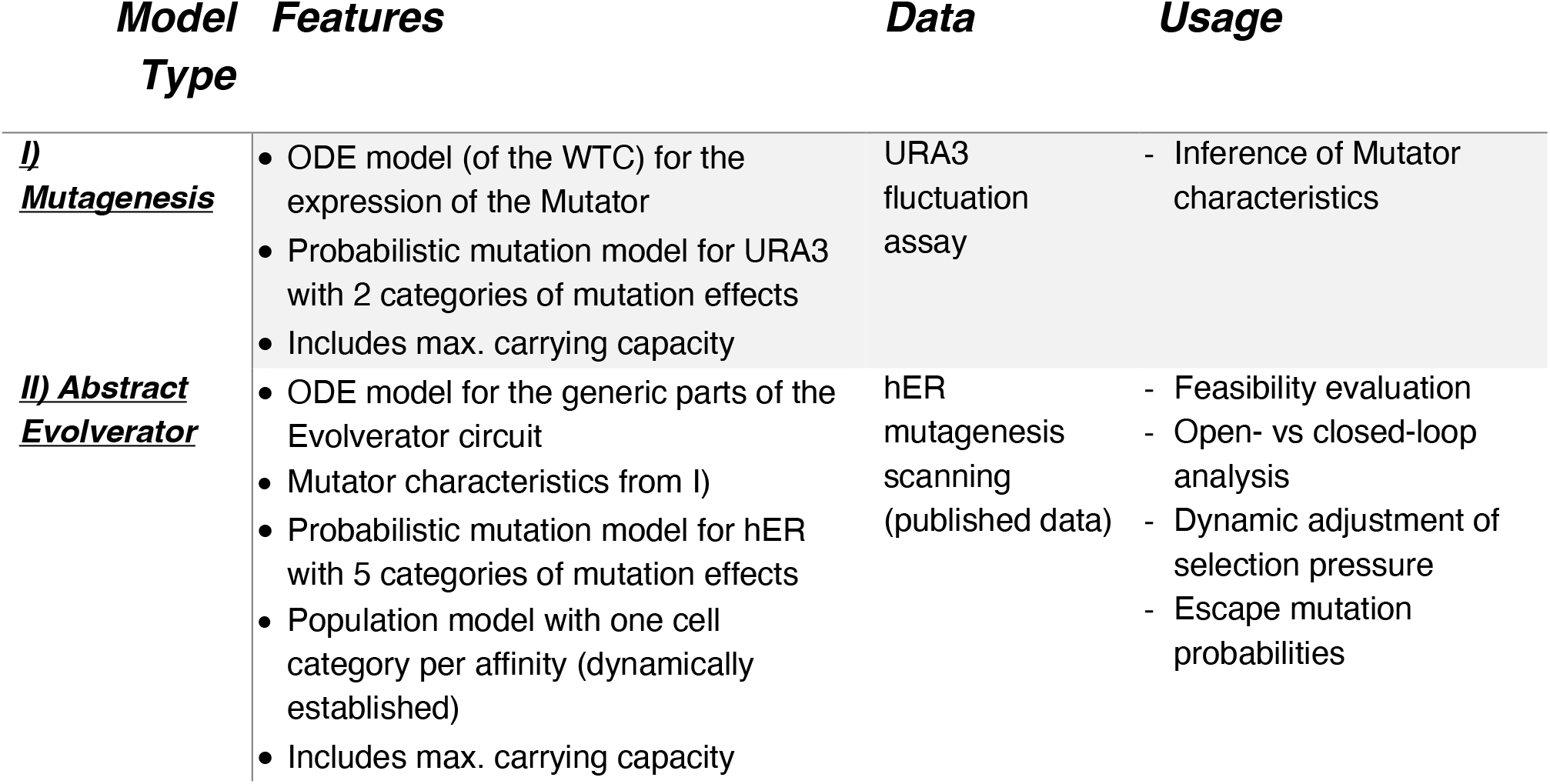
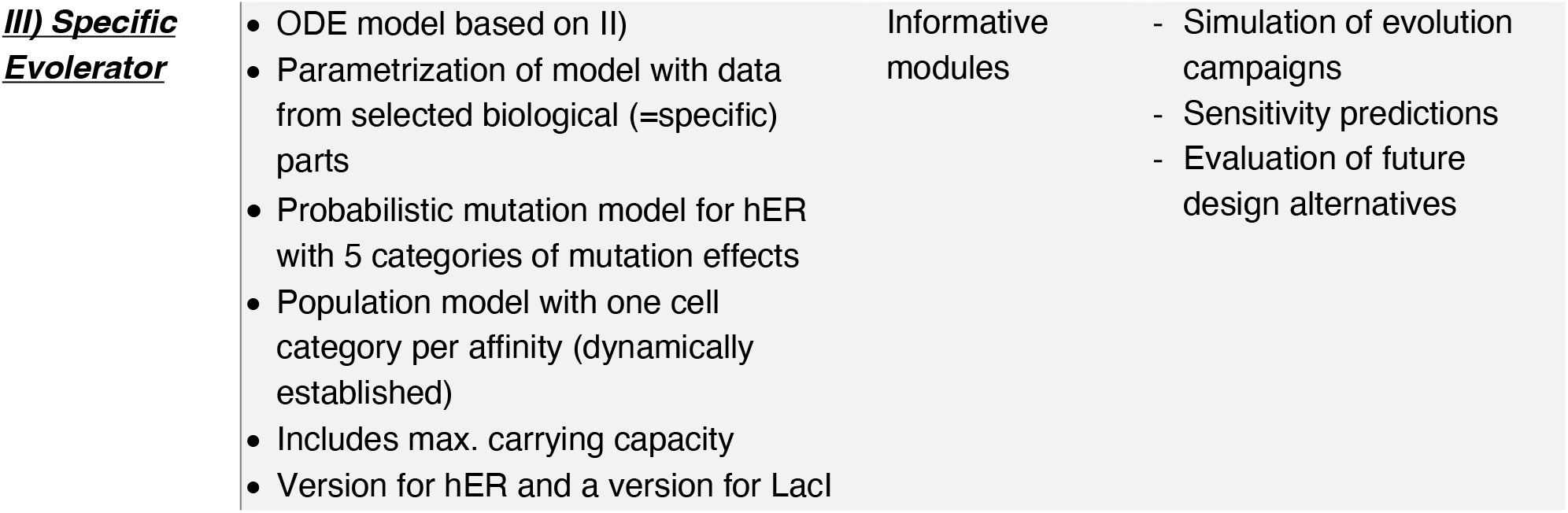

### Mutagenesis model for the URA3 gene

To capture the probabilistic process of accumulating mutations in a genetic sequence, we consider that in a given time-frame and for a fixed mutation rate, the numbers and positions of mutations (and therefore mutation effects) vary between individual cells. Our phenomenological model incorporates similar principles as prior work [48], namely that discrete groups of cells can arise due to mutations and that each group differs in their parametrization of the same ordinary differential equation (ODE) model. However, here, a mutator actively introduces mutations stochastically (not with a given mutation probability) and cultivation is in a batch culture, initiated with few cells and growing to saturation. We developed the model using preliminary fluctuation experiments with the MutaT7 system and the URA3 gene as screening marker[18] where expression of the T7 RNAP-cytidine deaminase was controlled by the WTC_846_ [14] (Figure S3B). The WTC_846_ is a genetic circuit where the gene of interest is expressed from a P7tet.1 promoter that is repressible by TetR, which is expressed from a P7tet.1 promoter as well (and represses its own synthesis). Basal expression is abolished by a second, constitutively expressed repressor: TetR-Tup1.

To model the concentration dynamics of the species of the WTC_846_ we adapted the published model [49]: TetR and TetR-Tup1 have their own Hill coefficients and we explicitly account for cell growth. For the total concentrations of TetR (_[*TetR*]_) and TetR-Tup1 (_[*TetRTup1*]_), the dynamics are:

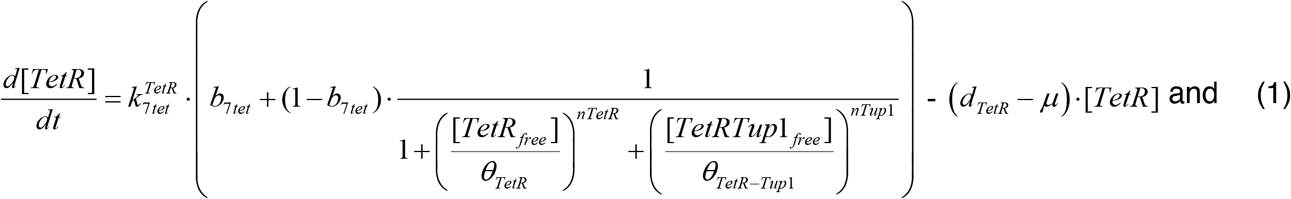

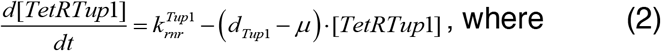

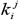 are maximal expression parameters of protein *j* from promoter *i, b*_*i*_ corresponding basal expression parameters, *θ*_*j*_ inhibition constants, *n*_*j*_ Hill coefficients, *d*_*j*_ degradation constants, and *μ* is the specific growth rate.

Assuming rapid equilibrium of the binding reactions between the repressors and the ligand aTc (_[*aTc*]_) [49], [50] as well as mass conservation, free and bound repressor concentrations are related by:

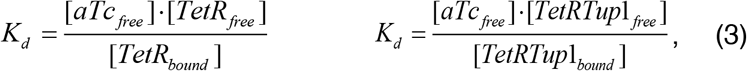

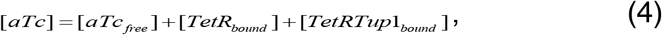

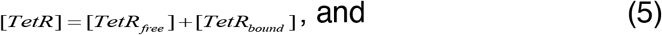

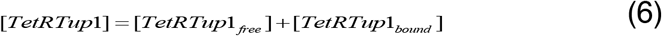

with the dissociation constant *k*_*d*_.

Solving for the active (free) TetR and TetR-Tup1 concentrations, we obtain:

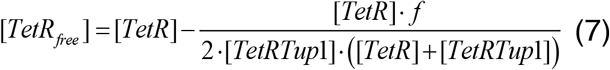

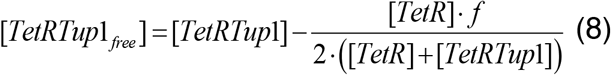

with

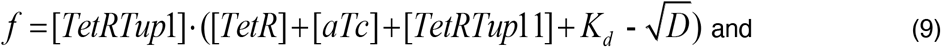

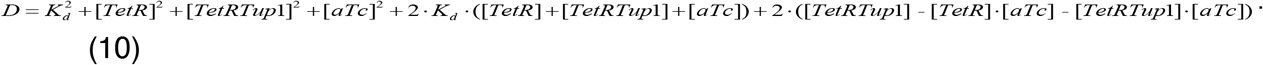

Instead of a fluorescent protein in the WTC_846_ [49], we model the dynamics of the mutator. While trying to fit our model to the time-course experiment (Figure 3B), we noticed an apparent time delay between the mutator being produced and the first cells with a mutation appearing. We therefore introduced inactive and active versions of the mutator ([*M*]),

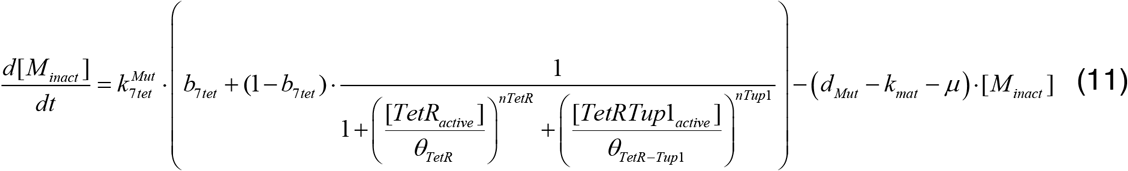

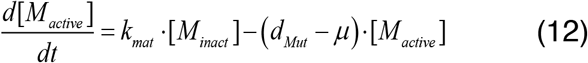

with maturation rate constant *k*_*mat*_. Moreover, mutator production decreases as cultures reach stationary phase (Figure S3C,D). We assume that gene expression capacity decreases in stationary phase and therefore, the production rate of each protein is set to almost zero when a culture reaches 90% of the maximal total cell density in stationary phase, *N*_*max*_.

Next, we model the mutation rate ρ for a given mutator concentration using a four-parameter logistic equation:

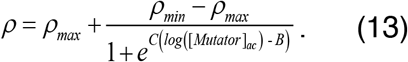

Here, ρ_*min*_ and ρ_*max*_ represent the mutation rate without mutator expression and upon maximal expression of the mutator from the P7tet promotor (obtained from the preliminary fluctuation assay), *B* presents the log EC50, and *C* the slope of the curve.

The specific growth rate of each cell was modeled by a logistic (Verhulst) equation with maximal specific growth rate ρ_*max*_ and carrying capacity *N*_*max*_

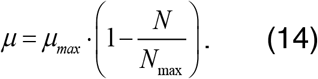

At the population level, we model two categories of cells: those with a deleterious mutation leading to a non-functional URA3 gene (cell number *N*_*LOF*_), and those without a mutation, or a mutation that keeps the URA3 gene functional (*N*_*F*_), leading to *N*=*N*_*LOF*_*+N*_*F*_. The number of cells in each group is given by the ODEs:

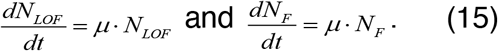

We model the number of mutations accumulating in the two subpopulations *k* within a time interval _[_*t*_*1*,_*t*_*2)*_ by drawing randomly from a Poisson distribution specified by the rate parameter λ [51], [52]:

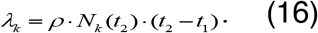

We expect that with our cytidine deaminase 8 out of 214 possible mutations result in a loss of function because they lead to a non-sense mutation [53]. For each mutation, we therefore decide if it is deleterious or not by randomly drawing the mutation type using the weights 8/214=0.0364 for a LOF mutation and 206/214=0.9626 for a mutation that keeps the URA3 gene functional.

We simulate the full model for the same duration as the experiment. After a time interval of 6 h, we predict the number of mutations that occurred, apply the mutations to the cells in the two groups and subsequently update cell numbers. As it is unlikely that an individual cell accumulates multiple mutations in 6 h, the number of mutations represents the number of cells that gained a mutation.

To parametrize the model, we used data from a fluctuation assay (Figure 3A, and from a separate time-course experiment (Figure 3B). We ran the model 25 times to resemble an experiment with 25 wells. For experimental and simulation results, we each fitted a log-normal distribution to the number of mutants. We then minimized the absolute difference of the estimated means using the enhanced scatter search method [54] (Figure S3E,F and Supplementary Table 2B).

### Evolverator model

To describe the dynamics of the Evolverator species (Target/Receptor, Inverter, Mutator, Growth protein) we formulate ODE systems similar to the mutagenesis model, with corresponding notation for model parameters (see also Supplementary Table 2C). As the exact promoter is not known yet, maximal expression parameters are given as *k*_*j*_ instead.

For the design with constitutively expressed receptor ([*R*]) (Figure 3D) we use:

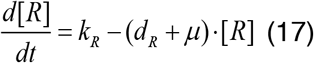

and for the design with negative feedback (Figure 3E) we use:

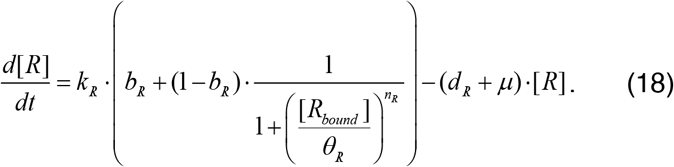

We model the expression of the inverter ([*I*]) and growth regulating protein ([*G*]) in a similar fashion:

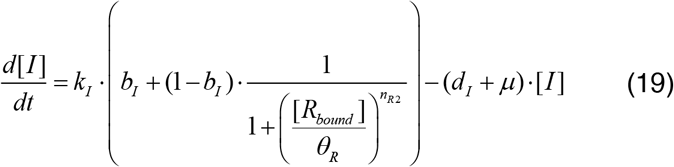

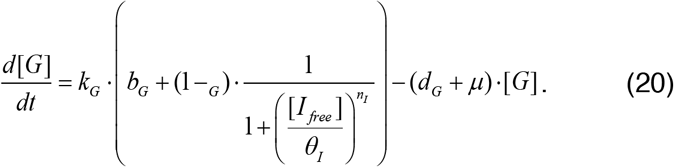

The ODEs for inactive and active mutator, analogous to Eqs. (11)–(12), are:

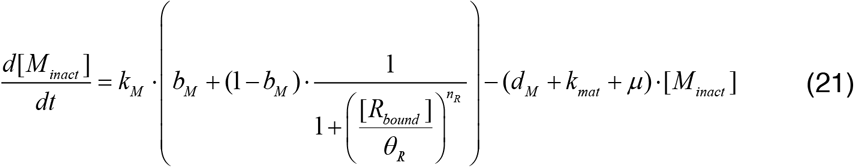

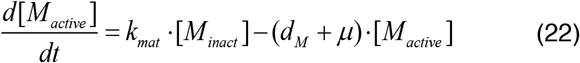

and the number of cells in a population is tracked by:

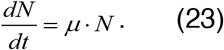

The receptor can only repress gene expression when bound to its ligand ([L]). Under the same quasi steady-state assumptions and exploiting mass conservation as for the mutator model, we define the amount of active receptor as:

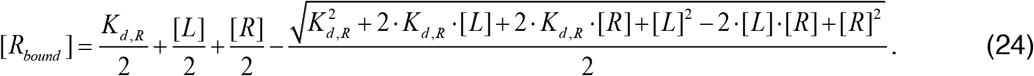

To fine-tune the expression levels of the growth protein, a second ligand (L_2_) can be added to tune the activity of the inverter. Here, we incorporate a mechanism in which the addition of a ligand inactivates the inverter(resembling the behavior of the repressor TetR), leading to:

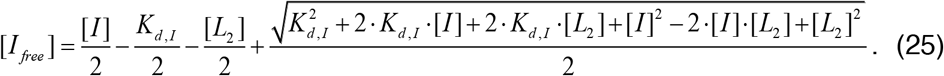

To map the mutator concentration to the mutation rate *ρ*, we use Eq. (13) and a similar four-parameter logistic equation to relate the expression level of the growth protein to the maximal growth rate, with parameters for the growth rate bounds indicated by asterisks:

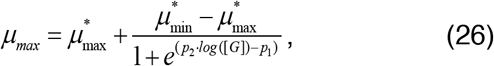

which then enters the logistic relation for culture growth, Eq. (14).

To capture mutagenesis and its effects on cell states and fitness, we model the introduction of mutations in an isogenic starting population. Hence, we start with a single group of cells from which discrete groups can arise that differ in their affinity to the ligand (see Figure 3C). Each discrete group of cells *k* has their own set of the model in Eqs. (17)–(26), but some of the parameters are adjusted to reflect the group’s associated affinity. Correspondingly, every time a new affinity appears in the culture, a new set of ODEs is added to the model for the new affinity group, and when an affinity is lost, the corresponding set of ODEs is removed. Formally, we indicate the group identity by expanding the notation above with subscript *k*. The total number of cells in a culture with *n* subgroups is then:

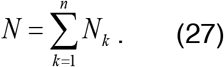

Note that all sets of ODEs for the different groups are simulated together, to model the dynamics of the whole population.

### Performance evaluation

To assess if an Evolverator design for a given model parametrization and given starting (*K* _*A,strart*_) and target (*K*_*A,target*_) affinities is predicted to be functional, namely to evolve suitable variants in a given time frame (Δ*t*), we evaluate three criteria (see also Figure 3D-E).

The first criterion, *C*_*µ*_, evaluates the affinity-growth rate relationship for discrete sets of log-transformed affinities **K**_**A**_ = {*K*_*A*,i_ ∈ [*K*_*A,start*,_ *K* _*A,target*_]}. For the design with feedback, aiming for a linear relationship between affinity and growth with zero growth at the starting affinity, maximum growth at the target affinity, and a single ligand concentration, we define:

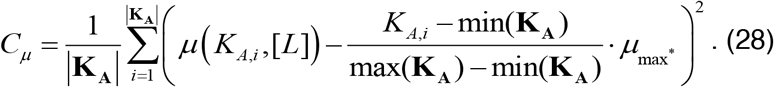

For the design without feedback, with sigmoidal dose-response curves that we shift by varying the ligand concentration, we require that for given affinities and matched ligand concentrations [*L*]_*i*_, the growth difference between the current and the next (intermediate) target affinity is a fraction *f*_*µ*_ of maximal growth:

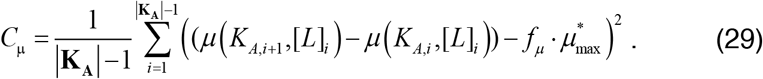

The second criterion, *C*_*ρ*_, tests if the starting mutagenesis rate is above a threshold (*ρ*_1_) to make sure enough mutations are introduced into the population and the mutagenesis system is being shut off (below a threshold *ρ*_2_) when the target affinity is reached:

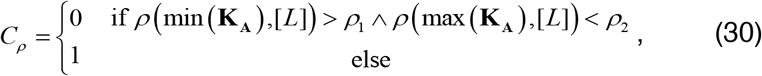

where different ligand concentrations may be applied for the design without feedback as above.

The final criterion, *c*_*evalution*_, assesses the change of affinity over time in the culture, specifically how many out of *n* wells reach our target affinity within the given time-frame. Denoting by 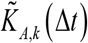 the dominant (i.e., highest frequency of corresponding cells) affinity over the cell groups in a given well *k* at the end of the evolution experiment and using the indicator function

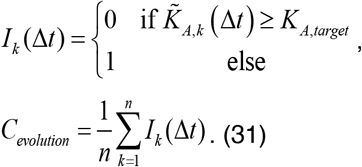

For computational efficiency, we evaluate the first two criteria by simulating the ODE system for different, fixed affinities (without introducing any mutations). If these criteria are met for a given parameter set, we run the model multiple times to represent an experiment with parallel cultivations in a 96-well plate.

More specifically, we started by setting a target affinity of 10 nM^-1^. We aimed for a 3-fold logarithmic change in affinity and therefore the starting affinity of the population was set to 0.01 nM^-1^. This represents a scenario where the target already has a decent affinity to the ligand, which we would like to further improve. Moreover, we define the target time as ~25 days, which is consistent with the period mentioned in previous continuous evolution experiments [26]. Maximal growth µ_max_ was set to 0.0077 min^-1^ (determined experimentally, see Figure S8A-B) and *f*_*µ*_ was set to 0.58. For the design without feedback, we fixed the starting ligand concentration to 25 nM. Subsequent ligand concentrations for shifting the growth-affinity curve were estimated together with the parameters. From our initial fluctuation assay (Figure 3A) we obtained minimum (for 0 ng/ml aTc) and maximum (for 600 ng/ml aTc) mutation rates of *2*.*10*^*−8*^ min^-1^ and *3*.*7*.*10*^*−6*^ min^-1^ (refer to ‘Fluctuation Assay’ above for calculation details). Therefore, ρ_1_ and ρ_2_ were set to *1*.*10*^*−6*^ and *5*.*10*^*−8*^ min^-1^ respectively. We only simulated the third criterion when *c*_*µ*_ < (0.0005)^2^ and *c*_*ρ*_ = 0.

### Adapting mutation probabilities to a receptor gene

To model feasibility of the Evolverator, we adapted the mutagenesis model’s mutation probabilities and effect from the URA3 gene to a receptor gene. We reasoned that for a receptor a mutation can lower, increase or not affect the affinity. Moreover, a directed evolution study of the human estrogen receptor[28] showed that mutations outside the ligand binding domain (LBD) affect the affinity less than mutations within the LBD. To parametrize the probabilities, we analyzed mutants from the same study in more depth. For each mutant, we took a codon representing the amino acid before evolution and determined the amino acids that can be obtained by mutating each nucleotide in this codon separately. When the mutation led to an amino acid within the same chemical class (based on their side chain (R group) type) to the amino acid that improved binding in the experiment, we classified it as beneficial. We defined neutral (same properties as the starting amino acid) and detrimental (different properties of the amino acid and no mutant with improved binding) mutations accordingly. We then averaged the results for the different mutants to obtain a probability for each mutation type. We also classified if a mutation was within or outside of the LBD. Additionally, we used a BLAST CD search with default parameters [55] to determine which amino acids were highly conserved, assuming that changing a conserved amino acid impacts the receptor in a detrimental way. Moreover, we used the results in [28], [56] to estimate the effect of the mutations on the affinity, resulting in the values compiled in Figure S3G.

The probabilities for beneficial mutations appear consistent with literature (0.01%-0.5%, [57]). However, to assess the sensitivity of our predictions to uncertainties in these estimates, we simulated our Evolverator model with different probabilities. Figure S3I shows that a change in the probability of a beneficial mutation has a large effect (leading to a different outcome), but changing the other mutation types only slightly affects the run time of an evolution experiment.

### Models for small, informative modules

To characterize the 18 small informative synthetic networks (called modules; see Figure S9 and Supplementary Table 2A) that we measured experimentally, we created a general model containing all components and interactions of all modules. The model of each individual module can be obtained from the general model by projecting the irrelevant parameters to zero, thereby eliminating the irrelevant reactions as in [50]. Parameter notation is as above, and for detailed definitions see Supplementary Table 2D.

The repressor dynamics are given by:

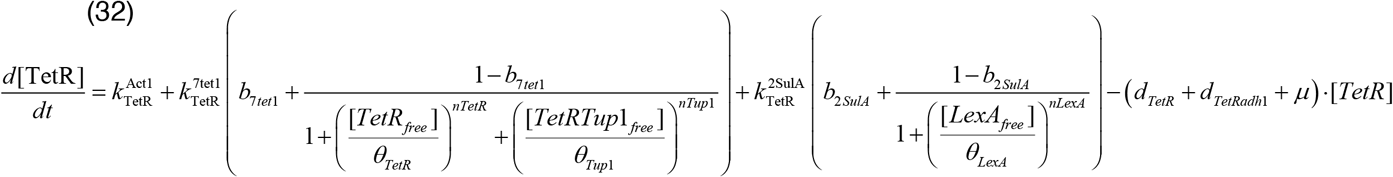

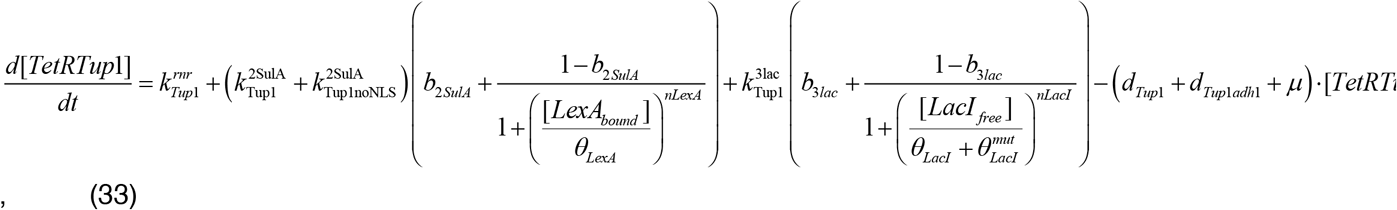

where we modeled the difference between TetR-Tup1 with and without the NLS tag by two different parameters for the production of TetR-Tup1, and correspondingly included two degradation parameters for protein variants with / without degradation tag.

The dynamics of PMA1, LexA (LexA-hER and LexA-hER-lac12 because they never appear in the same module), LacI (LacI and Citrine-LacI fusion protein, correspondingly), and the active / inactive mutator are captured by:

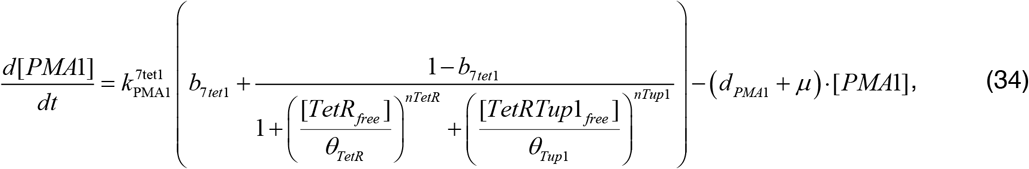

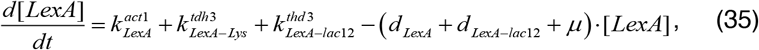

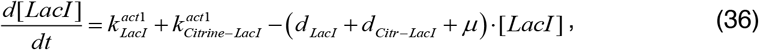

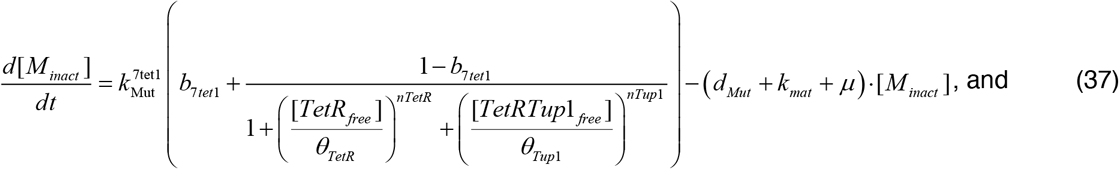

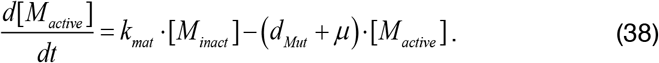

For the calculation of active protein forms, see the section Evolverator model; LacI_free_ was determined in the same way as TetR_free_. Moreover, we linked PMA1 concentration to growth rate via Eq. (26) and mutator concentration to mutation rate via Eq. (13).

We used the same ODEs for the two fluorescent proteins (FPs) mCitrine and mApple, accounting for protein maturation via two states:

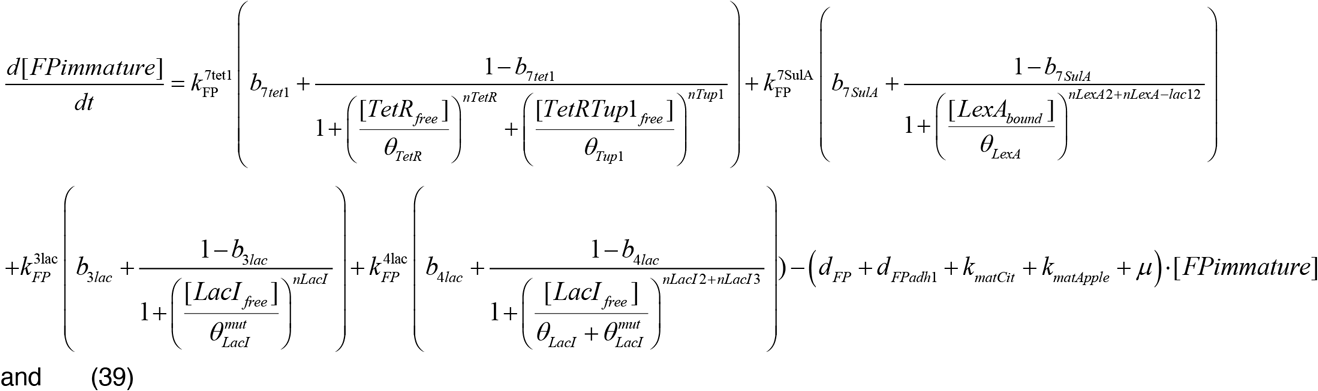

and

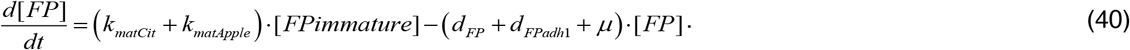

For the two FPs we assumed equal production and degradation rate constants as both are stable proteins and have a similar length. However, we allowed for different maturation rate constants *k*_*mat*_.

To relate the model simulations to fluorescence intensities obtained by flow cytometry in arbitrary units, we used scaling factors as in ref. [50]. For the growth data obtained by the Growth Profiler, we fitted a logistic growth equation (kinetic version) to the growth curves [58]:

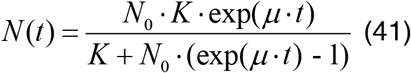

with carrying capacity K, an initial population size N_0_ and a specific growth rate *µ*.

Moreover, to check for effects of aTc and β-estradiol on cell growth and for modules with an assumed potential for creating burden for the cell because of the number of integrated components (e.g,, with fluorescence as output; see Supplementary Table 2A) we used the growth profiler and the same fitting procedure to estimate the growth rate of each strain in different experimental conditions (see Figure S8A,B). This growth rate was then assigned to the parameter *µ*.

### Estimation of mutation rates

We characterized our improved mutagenesis system as part of the Evolverator by performing a fluctuation assay for different levels of expressed mutators, after placing the mutator gene under the control of the WTC. This assay measures the occurrence of a loss-of-function (LOF) mutation in the URA3 gene (phenotypic mutation rate, per generation) and allows us to estimate the substitution per-base-pair (spbp) mutation rate of the combined deaminase pair. To estimate the spbp rate, the phenotypic mutation rate needs to be corrected with the effective target (bp) [53]. Commonly, one considers all the nonsense mutations that can be introduced with a single base pair substitution. However, we also included a set of missense mutations that we expect to lead to a LOF mutation (otherwise we could not calculate a rate for the abe8e system). These missense mutations were found by determining the conserved amino acids in the URA3 protein via a BLAST CD search with default parameters [55]. The estimated growth rate of our mutator strain for each aTc condition was taken into account to obtain a spbp rate in hours (instead of generations; see Figure S8C,D). The spbp mutation rate was used for fitting the parameters of the mutator. To obtain specific mutation rates for the hER and LacI genes for use in the model, we multiplied the spbp rate with the length of the hER and LacI gene (resp. 942, 1083 bp).

### Parameter Estimation

We used the experimental data for all modules jointly to estimate the unknown model parameters using the enhanced Scatter Search (eSS) algorithm of the Meigo toolbox [54]. After assembling the unknown parameters into a parameter vector Φ, we minimized the objective function (*χ*^2^ statistic):

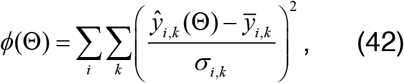

Where *ŷ*_*i,k*_(ŷ)represents the simulated fluorescence, mutation rate, or growth rate for dose *k* during experiment *i*, 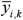 the mean of the corresponding experimental data, and the standard deviation of the experimental data. The optimal fit is shown in Figure S9 and the corresponding parameter values are given in Supplementary Table 2D.

### Simulating the evolution of hER and LacI

To predict the dynamics and outcome of evolution experiments, we simulated the Evolverator model,, using the parameter values estimated from the modules. We set the dilution times of the culture and the measurement times of the fluorescence to the experimentally used time points. We simulated the model 88 times, representing the 88 evolution wells. In addition, we simulated control wells once per control (note that the experimental results contain duplicates of the control wells). To obtain the correct starting affinity of hER for the simulations – as this parameter was not estimated for ligands other than β-estradiol – we simulated multiple dose response curves – differing by their K_d_ – and compared the EC_50_ value of each curve to the EC_50_ values of the dose response curves obtained experimentally (see Figure S10B). For each ligand, we chose the affinity with the closest match between simulated and experimental EC_50_.

To compare model outputs to experimental results, we converted the model’s cell numbers to cell density using a calibration experiment (see Figure S10A). To summarize the growth curve matches in a more quantitative way, we first fitted a non-linear mixed effect model to each growth period (with the cell density _Y_ in log-scale) for simulations and experiments:

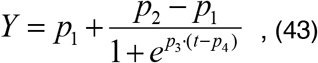

Where p_1_ and p_2_ are the cell densities of an empty and saturated culture, p_3_ the slope and p_4_ the mid-point (the times at which we reach the inflection point of the growth curve). Each parameter was assumed to be a sum of a fixed and a random effect. Covariances were not significant and therefore set to zero). We then compared estimated mid-points (see Figure 6C, Figure S10C-E).

### Model sensitivity

We analyzed how sensitive outcomes of evolution experiments are to changes in parameters by taking the parametrized Evolverator model and systematically changing the value of each parameter individually, while keeping the other parameters constant, and recording the number of successfully evolved wells as outcome (Figure 6D). Note that, during these simulations, wells were diluted upon saturation.

### Parameter sampling

We used the PESTO toolbox [59] to sample parameter values via Markov chain Monte-Carlo (MCMC) sampling to characterize the parameter space that can result in a working system and to identify potential parameter correlations (which need to be considered when adapting the system to different targets). We maximized the following likelihood by sampling:

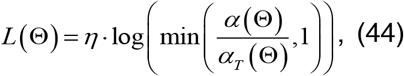

where *α(Φ)*is the sum of the affinities reached in each well, *α(Φ)*_*T*_the sum of all affinities when each well reaches the target affinity, and *η* a user-set parameter that defines the “steepness” of the log-likelihood curve. Again, we aimed for a three-fold log change in affinity and our starting affinity was set to the starting affinity of hER to estriol. In the corresponding simulations, we diluted each well upon saturation.

### Evaluation of design alternatives

To check if our system can be (easily) adapted to other evolution scenarios or targets, we performed forward simulations. For the scenario where we want to evolve a repressor with a ‘weak’ starting affinity, we took our parameterized evolution model for LacI, lowered its repression coefficient by a 100-fold and tuned the maximum expression rate of the repressor to obtain a working evolution system; this achieved a one-fold log change in affinity. To adjust selection pressure and allow for a three-fold log change in repression efficiency, we additionally tuned the concentration of a small molecule that modulates the concentration of active repressor.

For the second scenario we aimed to evolve protein-protein interactions. We replaced the ODE model of the target by the two ODEs:

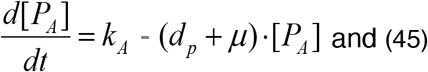

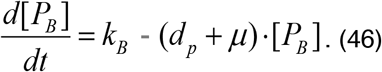

Under the same quasi steady-state assumptions as before and exploiting mass conservation, we define the concentration of protein-protein complex as:

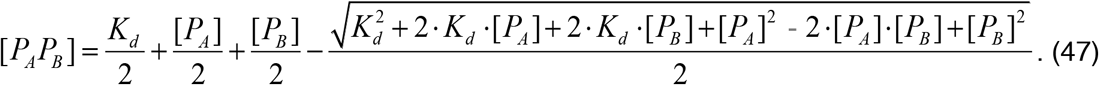

The protein complex [*P*_*A*_*P*_*B*_]then acts on the inverter and mutator modules as a repressor. We kept the equations and parametrizations of the mutator, inverter and growth modules as before. For [*P*_*A*_]and [*P*_*B*_], we then tuned the parameters to obtain a working system, staying close to the previously estimated values of the lexA-estrogen receptor to make sure the parameters are realistic. To change selective pressure, the parameter k_A_ was tuned with each log fold change in affinity.

