## Supplementary Figures for "Evolverator: An engineered yeast system to drive rapid continuous *in vivo* evolution of proteins"

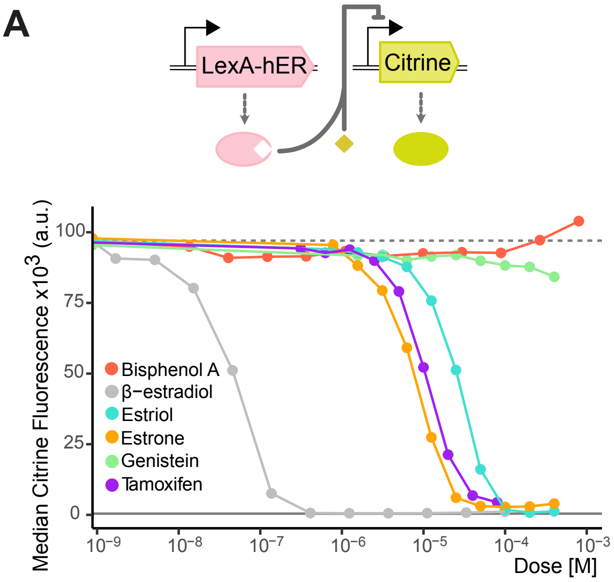


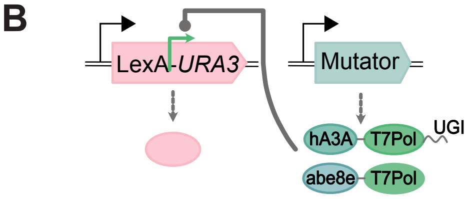


| *Mutator* | none | hA3A | abe8e | both |
|---|---|---|---|---|
| *Mutation rate (x10$^{-7}$)* | 0.44 | 129 | 17.2 | 187 |
| *95% Confidence Interval (x10$^{-7}$)* | 0.21-0.75 | 102-155 | 13.2-21.4 | 157-216 |
| *Fold change* | 1 | 293 | 39 | 425 |

**Supplementary Figure1. Molecular components of the Evolverator. A)** LexA-hERLBD displays different affinities towards different hormone ligands. Top: Strain where $P_{7SulA.1}$ controls Citrine expression and $P_{ACT1}$ controls LexA-hERLBD expression (Y3516). Diamond = ligand. Bottom: Dose response of Y3516. Dots: median Citrine fluorescence from flow cytometry (n=5000 cells per dose); solid horizontal line: median autofluorescence (Y70); dashed horizontal line: median maximum $P_{7SulA.1}$ activity (Y3391). **B)** Mutation rates of mutators in a fluctuation assay designed to deactivate Ura3. Top: Schematic of the target ($P_{ACT1}$_LexA-$P_{T7}$-*URA3*) and the mutators inducibly expressed by $WTC_{846}$. Green arrow: T7 promoter. Bottom: The four strains tested (Y3119, 3858, 3859, 3860) had the indicated mutators genomically integrated and no other genomic copy of *URA3.* The number of 5'FOA-resistant colonies was used to estimate the number of mutations acquired per cell per generation (see STAR Methods).

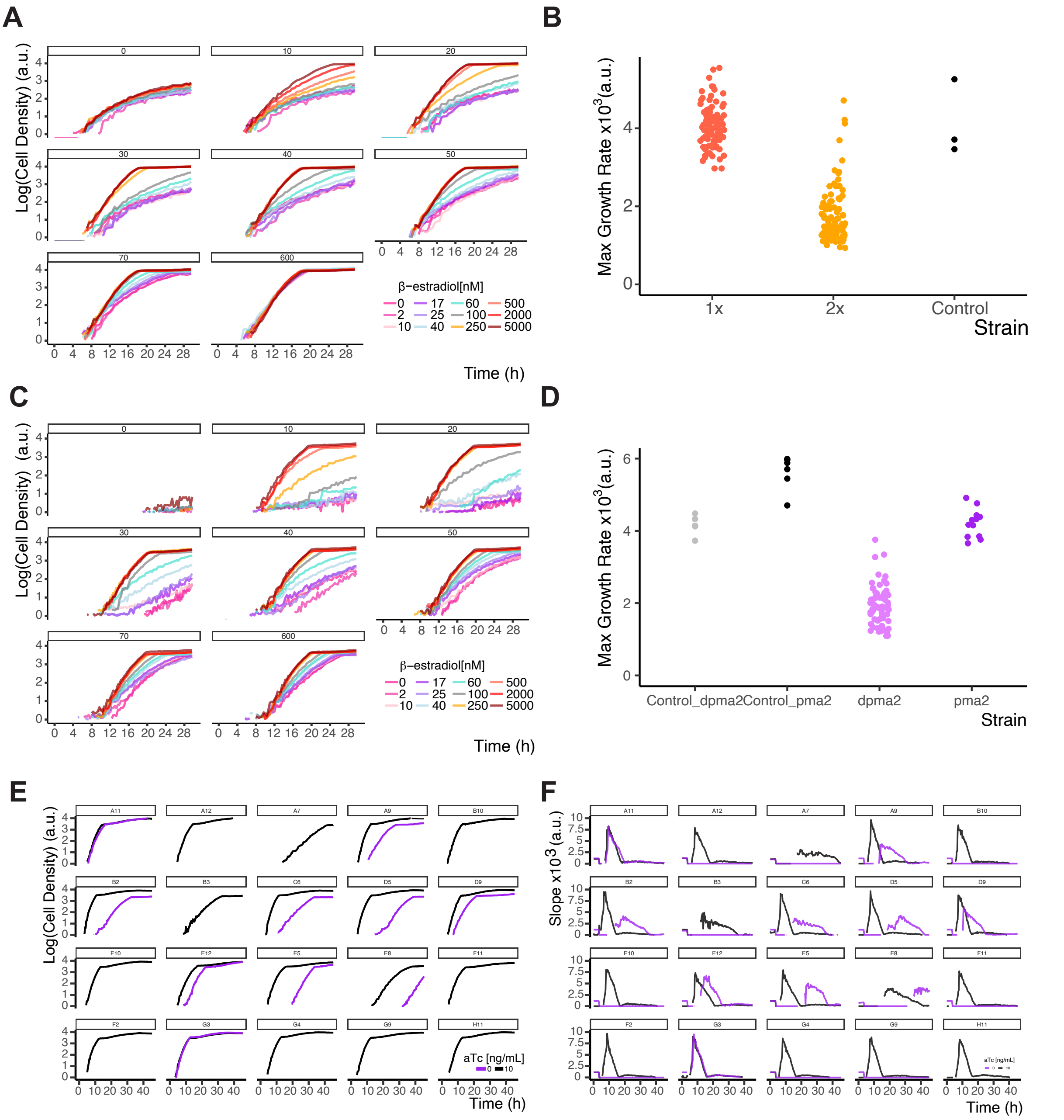


**Supplementary Figure 2. Growth control through *PMA1* and the effect of escape mutations. (A)** A strain in which $P_{ACT1}$_LexA-hERLBD repressed $P_{2SulA}$_TetR-Tup1, which in turn repressed $P_{7tet.1}$_*PMA1* (Y3435, same as Y3883 but without the mutators), was grown in liquid YPD with varying β-estradiol and aTc concentrations, and culture density was recorded continuously. Numbers above the plots indicate aTc concentrations in ng/mL. (**B)** Maximum slopes observed in the growth curves in **Figure 2A** (**STAR Methods**). **(C)** A strain in which $P_{ACT1}$_LexA-hERLBD repressed two copies of $P_{2SulA}$_TetR-Tup1, which in turn repressed $P_{7tet.1}$_*PMA1* (Y3884), was grown and measured as in (A). (**D)** Same as in (B) but for data in **Figure 2B**. (**E)** Wells of the 2x strain (Y3884) that showed increased growth in the experiment in **Figure 2A** were diluted at a ratio of 1:250 into YPD with 10ng/mL aTc or without aTc, to determine which component of the Evolverator was likely to have been mutated. Growth curves as in (A). Growth only in presence of aTc indicates a likely escape mutation at the target gene LexA-hERLBD. Fast growth in both conditions indicates that both TetR-Tup1-AT copies were likely deactivated. Fast growth with aTc, coupled with slower growth without aTc, indicates a likely mutation at only one copy of TetR-Tup1-AT. (**F**) Slopes of the growth curves presented in (E).

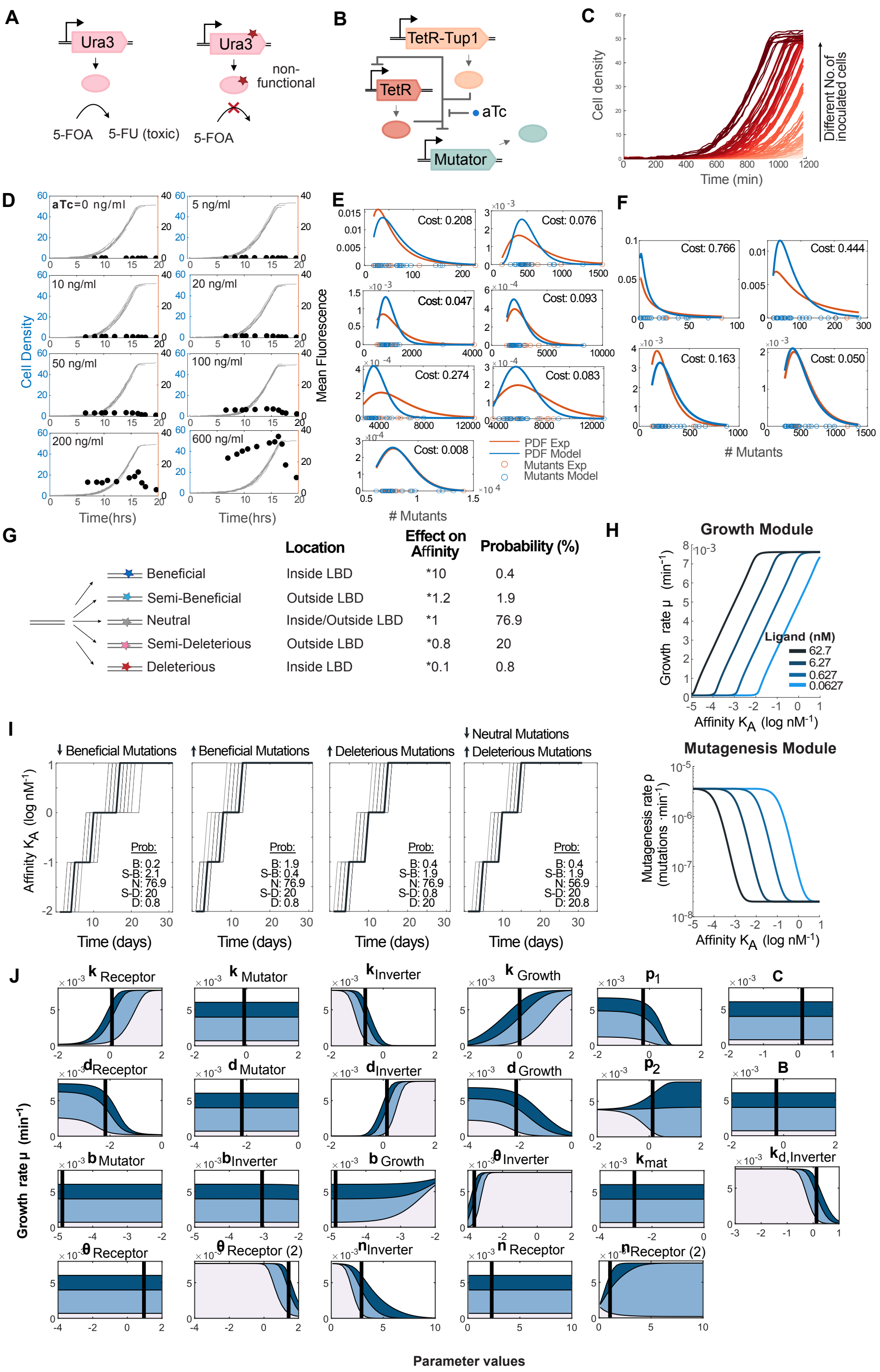


**Supplementary Figure 3. Development of the Mutagenesis and Evolverator model.**
**(A)** A fluctuation assay was performed using the URA3 gene and 5’ FOA selection. The Ura3 protein converts 5’FOA to a toxic intermediate, and therefore only cells where one or more mutations deactivated Ura3 can form colonies on the 5’FOA plates. **(B)** Schematic of the $WTC_{846}$ controlling the

expression of the mutator: a T7 polymerase fused to a cytidine deaminase. **(C)** Cells bearing the circuit from (B) - with additionally a Citrine fluorescent protein attached to the mutator- were grown in liquid YPD with varying starting numbers of cells (represented by the different colors) and aTc concentrations. Cell density (a.u.) was recorded continuously with a Growth Profiler 360 by taking images of the plate every 30 minutes. **(D)** Fluorescence was measured at different stages in the growth curve in (C) using flow cytometry. For each aTc concentration, the growth curves arising from the different starting conditions were shifted on top of each other (grey lines, left y-axis) and the corresponding fluorescent measurements (black dots, right y-axis) were then plotted as a pseudo time course. (**E, F)** A log-normal distribution was fitted to the number of mutants (line: fit, dots: experimental data) for each aTc condition and time-point measurement in the fluctuation assays (see **Figure 3A,B**). The score in each plot represents the difference in mean of the fitted distributions between the experimental (orange) and simulated (blue) results. (**G)** Type and frequency of mutations that can occur in our evolution model for hER and their corresponding effect on the affinity. For example, when a cell has an affinity of 1 nM and obtains a beneficial mutation, its affinity becomes 10 nM. (**H)** Model A (**Figure 3D**) can be adapted to different starting affinities by changing the ligand concentration**. (I)** Effect of modifying the probability of different mutation types on the functioning of the Evolverator (Model B, see **Figure 3E**). (**J**) The effect of changing parameter values – one at a time – on the growth rate obtained during evolution experiments for different starting affinities (different shades of blue). Parameter meanings are explained in STAR Methods and Supplementary Table 2C. The black line in each panel indicates the parameter value obtained from parameter estimation on small, experimental modules (see Supplementary Table 2D) that was used during the simulations.

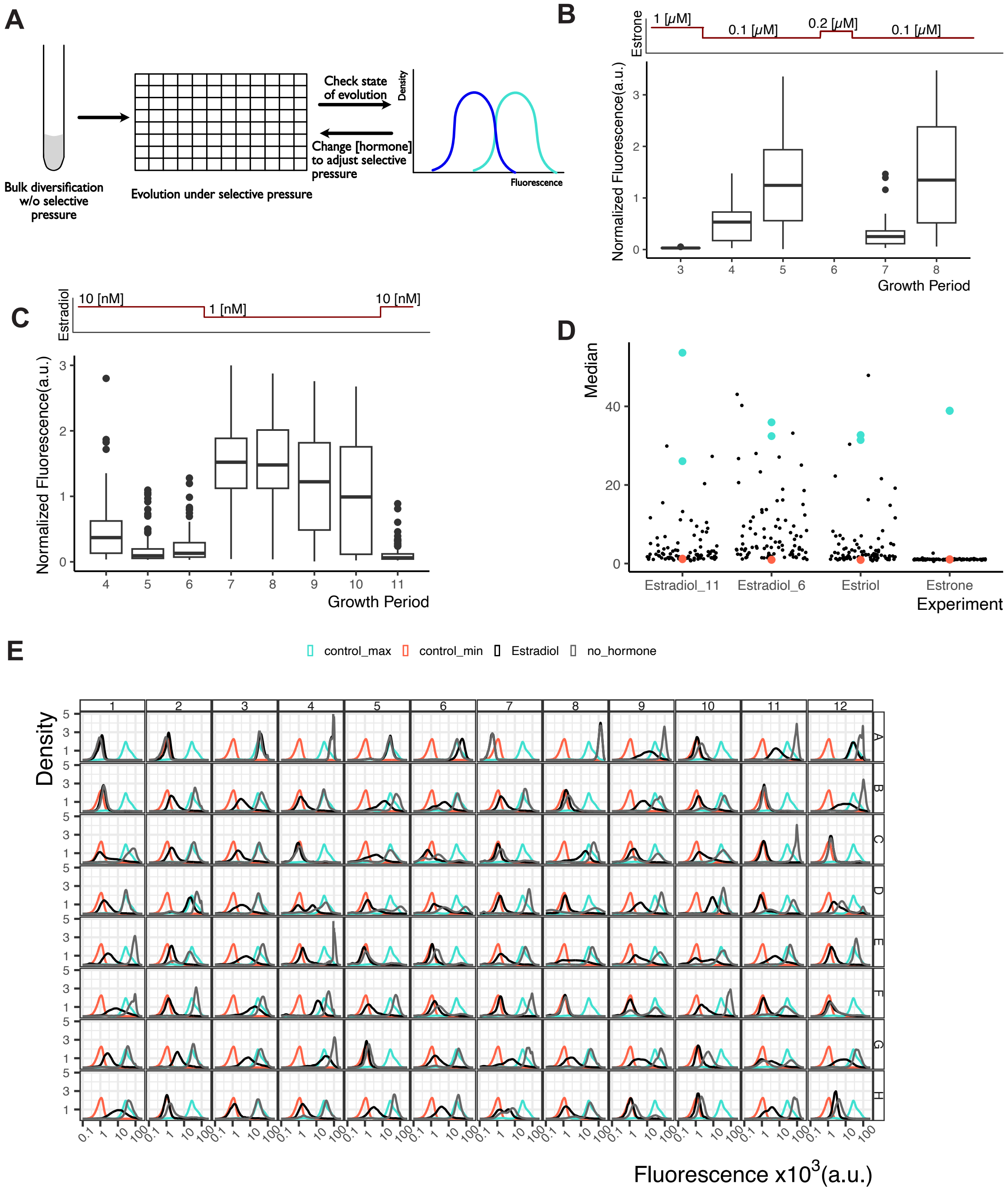


**Supplementary Figure 4. Tracking the state of evolution using a fluorescent reporter. (A)** State of evolution throughout the estrone evolution experiment with the Evolverator strain (Y4037). Citrine fluorescence was measured in each well using flow cytometry and normalized to the control well without hormone. Values below 1 indicate increased repression by LexA-hERLBD. Boxplots indicate median (line) and $25^{th}$ and $75^{th}$ percentiles (box). Estrone concentrations were as follows during each growth period. 1&3: 1μM, 2: 0.5μM, 4,5,7,8: 0.1μM and 6: 0.2μM. **(B)** Same as in (A), but for the β-estradiol evolution experiment. β-estradiol concentrations were as follows during each growth period. 1-6,11: 10nM and 7-10: 1nM. (**C)** State of evolution in each well at the end of the individual evolution experiments with the three hormones. Data shown in (A) and (B) were plotted as individual data points, each corresponding to one evolving well. Turquoise: control wells with 600ng/mL aTc and no hormone, red: control wells with 600ng/mL aTc and 10μM β-estradiol. (**D)** Wells A9 through to H12 from the β-estradiol evolution experiment, growth period number 11 were diluted into YPD with 4ng/mL aTc and either no hormone (grey) or 10μM β-estradiol (black) and grown to exponential phase. The control wells

A1&A2 had 600ng/mL aTc and 10µM β-estradiol, A3&A4 4ng/mL aTc, A5&A6 600ng/mL aTc and wells A7 (autofluorescence, Y70) as well as A8 (max fluorescence, Y3668) were internal controls for flow cytometry (grown in only YPD). Each well was sampled, Citrine fluorescence was measured with flow cytometry and plotted as density plots. Red: well A1, turquoise: well A6.

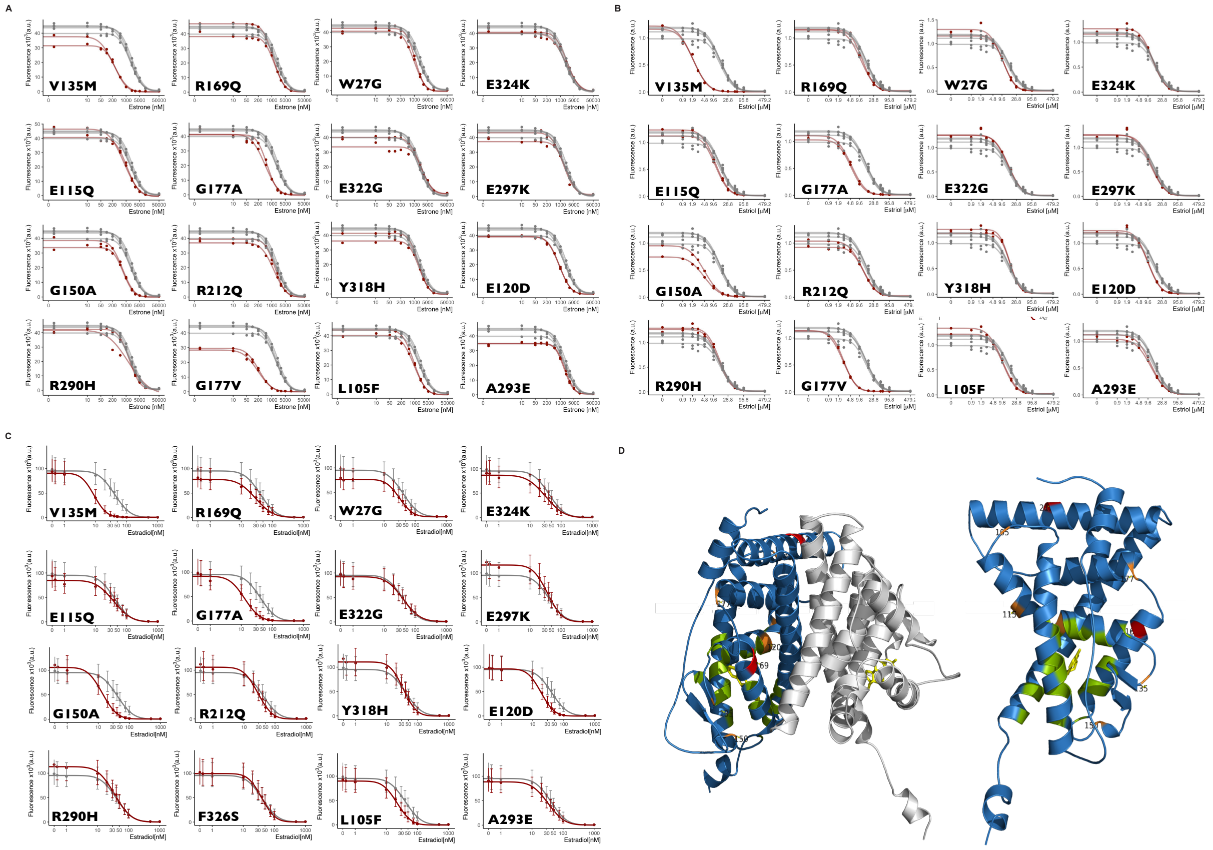


**Supplementary Figure 5. Characterization of single hERLBD mutations.** Single mutations found in the evolution experiments were cloned into a plasmid bearing $P_{ACT1}$_LexA-hERLBD and genomically integrated into a strain bearing $P_{7SulA.1}$_Citrine (Y4045,4046,4047,4052,4056,4057,4067,4069,4072,4073,4074,4075,4076,4078,4079,4081). These strains were grown overnight in different (**A)** estrone, **B)** estriol, and **(C)** β-estradiol concentrations and diluted into the same conditions. Citrine fluorescence was measured with flow cytometry 7 hours after dilution. Grey curves: strain bearing unmutated LexA-hERLBD (Y4045), red curves: strain bearing the mutation indicated at the bottom left corner of the plot. Dots indicate median fluorescence of the population and error bars span $25^{th}$ to $75^{th}$ percentiles. Dose response curves were fitted as explained in STAR Methods. **D)** Crystal structure of the hERLBD homodimer (left) and a single monomer (right) (PDB ID: 1A52) in complex with β-estradiol (yellow). In one monomer, the canonical binding pocket residues are colored green, and the mutations found in this work are colored red (mutations that affect basal activity) and orange (mutations that cause a left-shift in the dose response curve). Mutated residue numbers are displayed.

**A**

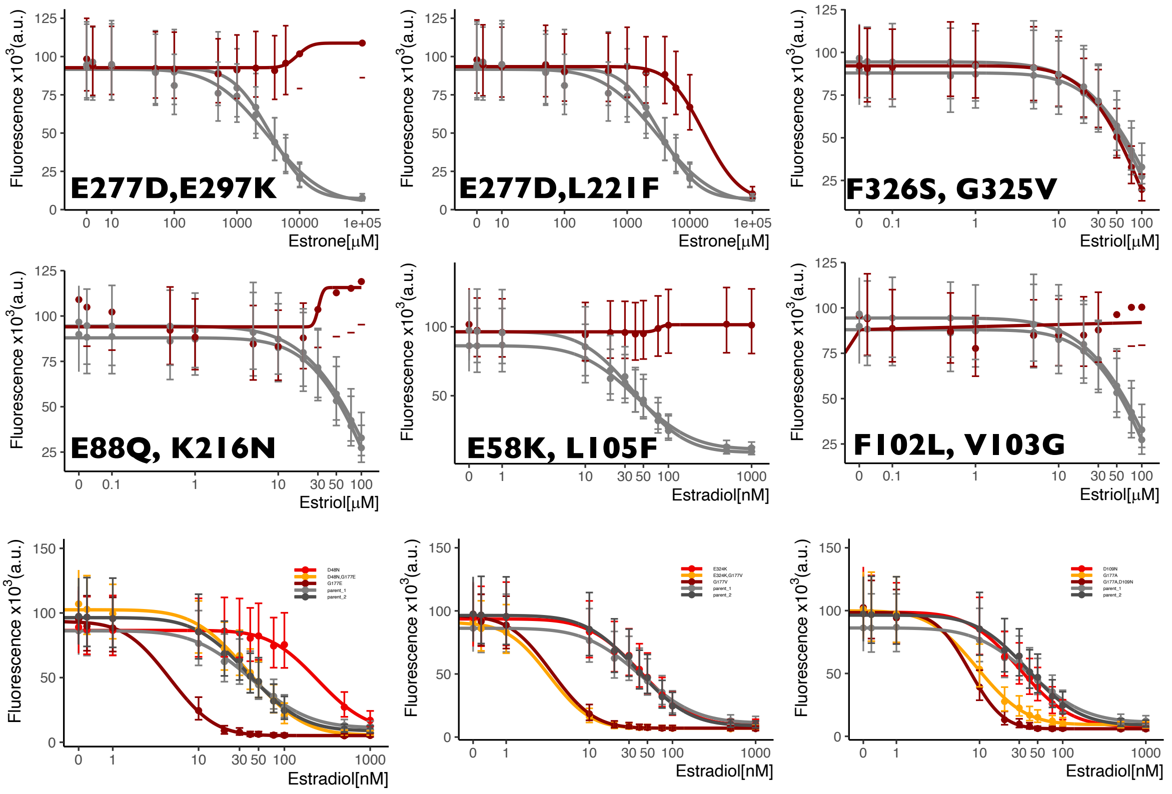


**B**

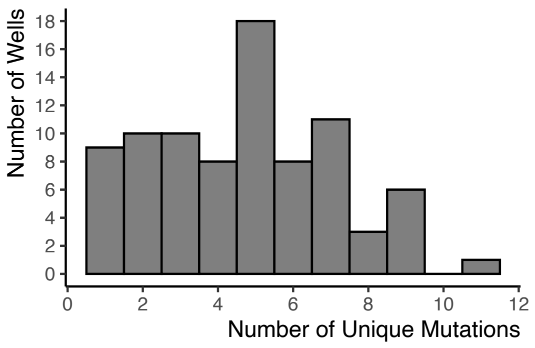


**C**

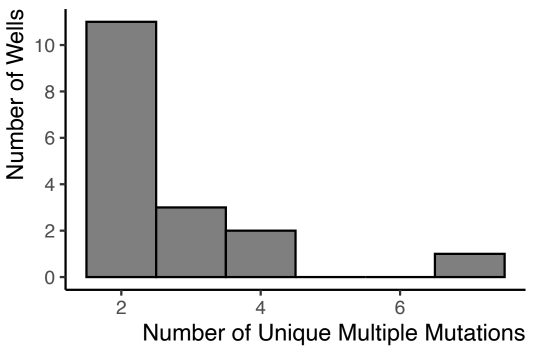


**Supplementary Figure 6. Characterization and prevalence of hERLBD mutations.** (**A)** Double mutations found in the evolution experiments were cloned as in Supplementary Figure 5A (Y4087,4089,4093,4094,4095,4096,4097,4098). These strains were grown overnight in different hormone concentrations and diluted into the same conditions. Citrine fluorescence was measured with flow cytometry 7 hours after dilution. In plots without a legend, grey curves indicate the parent strain bearing unmutated LexA-hERLBD (Y4045) and red curves indicate the strain bearing the mutation shown at the bottom left corner of the plot. Dots indicate median fluorescence of the population and error bars span 25$^{th}$ to 75$^{th}$ percentiles. Dose response curves were fitted as explained in STAR Methods. (**B,C)** Histograms showing the number of single, unique mutations (B) and of unique, multiple mutations (C) found per well at the end of the β-estradiol evolution experiment.

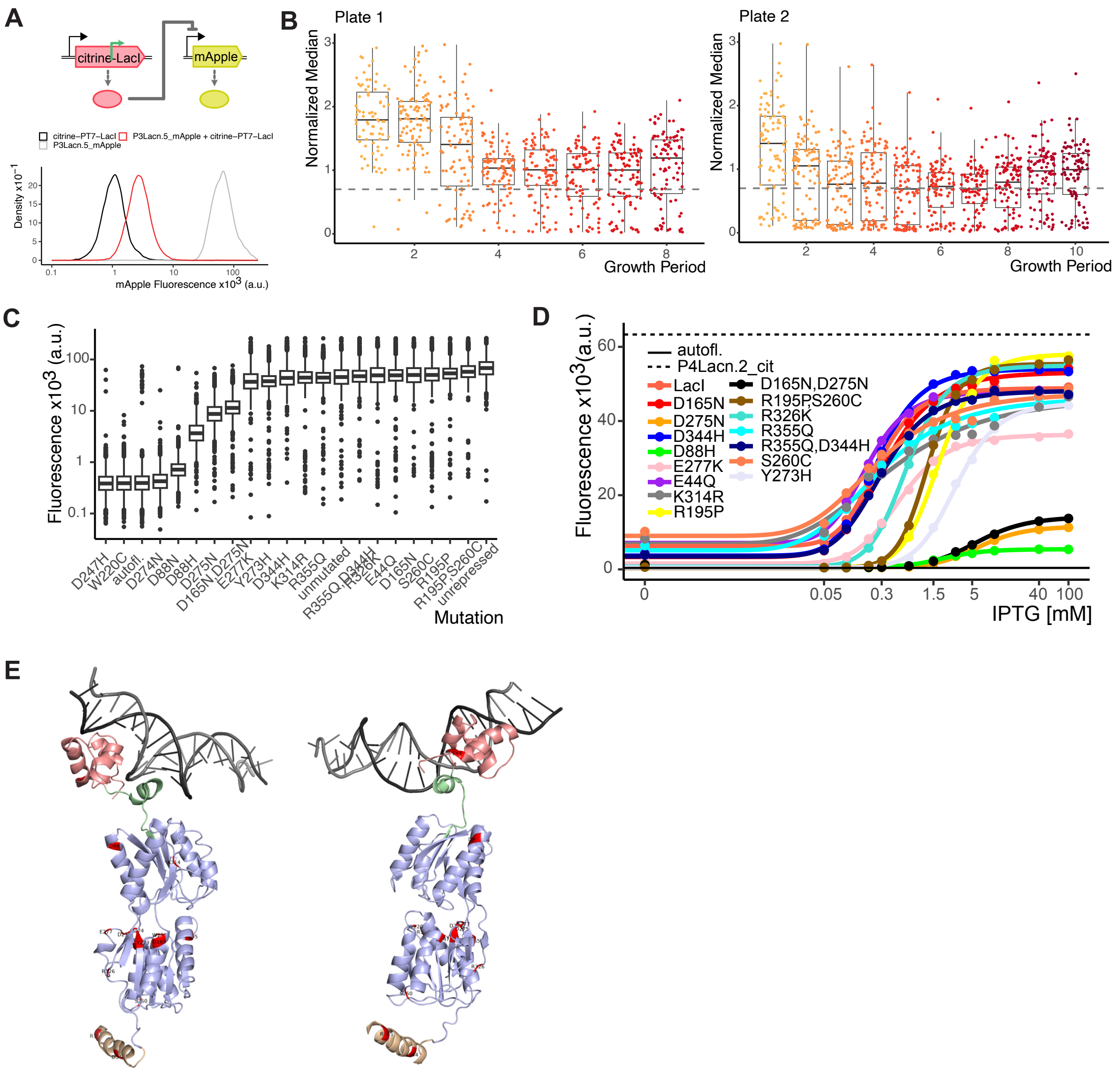


**Supplementary Figure 7. Evolving LacI with the Evolverator and characterization of the mutations found. (A)** Schematic representation of the strain Y4004, where the fluorescent protein mApple is expressed from the LacI repressible promoter $P_{3Lacn.5}$. The fusion protein citrine-$P_{T7}$-LacI is expressed from the endogenous yeast promoter $P_{ACT1}$. Dashed lines: transcription and translation, green arrow: T7 promoter. This strain was grown to exponential phase in YPD and mApple fluorescence was measured using flow cytometry. This was compared to mApple fluorescence measured in a strain that has only the $P_{ACT1}$ expressed citrine-$P_{T7}$-LacI (Y4003), and a strain where $P_{3Lacn.5}$ expresses mApple without any LacI (Y3729). Fluorescence values of the entire cell populations were plotted such that the area under the curve is one. (**B)** The Evolverator strain (Y3869) was grown overnight in presence of aTc (for unrestricted growth) and β-estradiol (for suppressing mutagenesis) and diluted into two separate 96 well plates in YPD at a concentration of 1000 cells per well. Plate 1 contained only rich media and Plate 2 contained 4ng/mL aTc. At the start of the 6$^{th}$ growth period, Plate 2 was also switched to the same conditions as Plate 1. During each growth period, samples were taken and mApple fluorescence was measured in each well. Fluorescence was normalized to the average fluorescence of the four control wells growing with 600ng/mL aTc and 10μM β-estradiol. Each dot indicates median fluorescence of a single well, colors indicate the growth period. The boxplot spans 25$^{th}$ to 75$^{th}$ percentiles and the line indicates the median of all evolution wells. The dashed horizontal line indicates 0.7-fold reduced fluorescence compared to control. (**C)** 16 unique single and 3 unique double mutations found during the LacI evolution experiments tested in Figure 5 (Y3965,3968:3972,3974,3976,3977,3982:3991) were grown in rich media with 20mM IPTG and Citrine fluorescence was measured using flow cytometry. Autofluorescence refers to the parent strain with only the Lac12 construct integrated (Y3316). Unrepressed refers to a strain where only the Lac12 and $P_{4Lacn.2}$ expressed Citrine constructs were integrated (Y3542). Unmutated refers to the original LacI sequence. (**D)** Selected strains were grown with various IPTG concentrations overnight and diluted into the same conditions to generate a full dose response. Citrine fluorescence was measured with flow cytometry. Solid line indicates autofluorescence (Y3316) and dashed line indicates maximum expression (Y3542). Dots indicate the median fluorescence of the population. Dose response curves were fitted using a 5-parameter log-logistic function (see STAR Methods). **E)** Crystal structure of a LacI tetramer bound to DNA (PDB: 1LBG) was modified so that only a single LacI monomer and the DNA operator are visible. Two separate views are displayed. DNA is gray. Different domains of LacI are indicated by colors: DNA binding domain is pink, hinge region is green, the core region is blue and the tetramerization domain is beige. Validated mutations found by Evolverator are indicated in red.

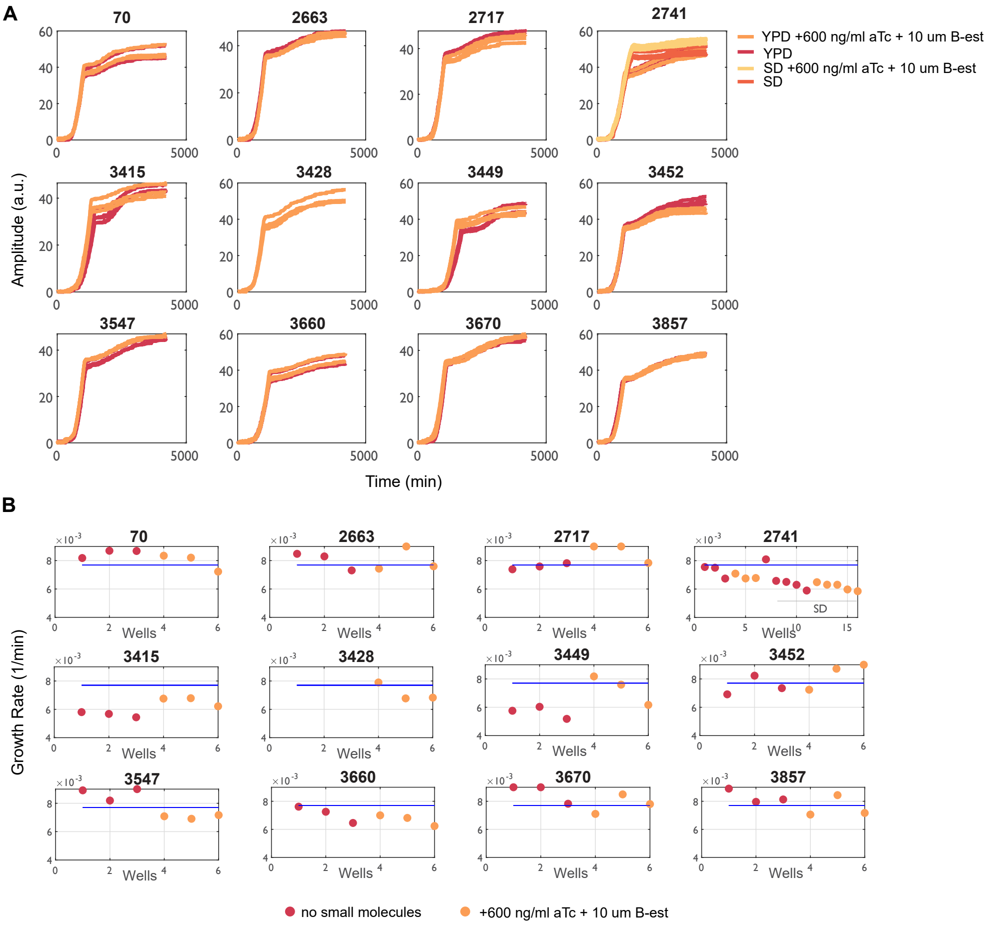


C

| aTc (ng/ml) | Phenotypical Rate (per gen) | pbp Substitution rate (per gen) |
|---|---|---|
| 0 | $2.45e^{-07}$ ($1.62e^{-07}$, $3.43e^{-07}$) | $3.27e^{-09}$ ($2.16e^{-09}$, $4.57e^{-09}$) |
| 50 | $1.39e^{-05}$ ($1.17e^{-05}$, $1.60e^{-05}$) | $1.85e^{-07}$ ($1.56e^{-07}$, $2.13e^{-07}$) |
| 100 | $2.56e^{-05}$ ($2.15e^{-05}$, $2.96e^{-05}$) | $3.41e^{-07}$ ($2.87e^{-07}$, $3.95e^{-07}$) |
| 200 | $3.64e^{-05}$ ($3.08e^{-05}$, $4.18e^{-05}$) | $4.85e^{-07}$ ($4.11e^{-07}$, $5.57e^{-07}$) |
| 600 | $4.79e^{-05}$ ($3.91e^{-05}$, $5.66e^{-05}$) | $6.39e^{-07}$ ($5.21e^{-07}$, $7.55e^{-07}$) |

D

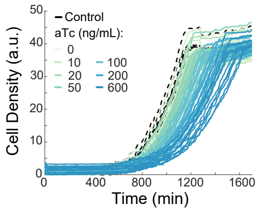


**Supplementary Figure 8: Estimating growth and mutagenesis rates from a selection of strains used for the parametrization of the Evolverator model.**

**(A)** Growth curves obtained for 12 different strains (top: strain numbers) grown in YPD with (orange curves) and without (red curves) aTc (600 ng/mL) and β-estradiol (10 µM). Strain 2741 was grown in SD media as well (yellow). For a description of each strain, see Supplementary Table 2A. **(B)** Estimated growth rates from the growth curves in (A) obtained by fitting a kinetic, logistic model to each curve (see STAR Methods). Blue line represents a division time of 90 minutes. **(C)** Per base-pair (pbp) substitution rate obtained from a fluctuation assay with different expression levels of mutator (see STAR Methods for the calculation). (D) The growth rate of the strain under different experimental conditions (aTc concentrations) was recorded and used in (C) to obtain a pbp rate per generation (instead of per hour).

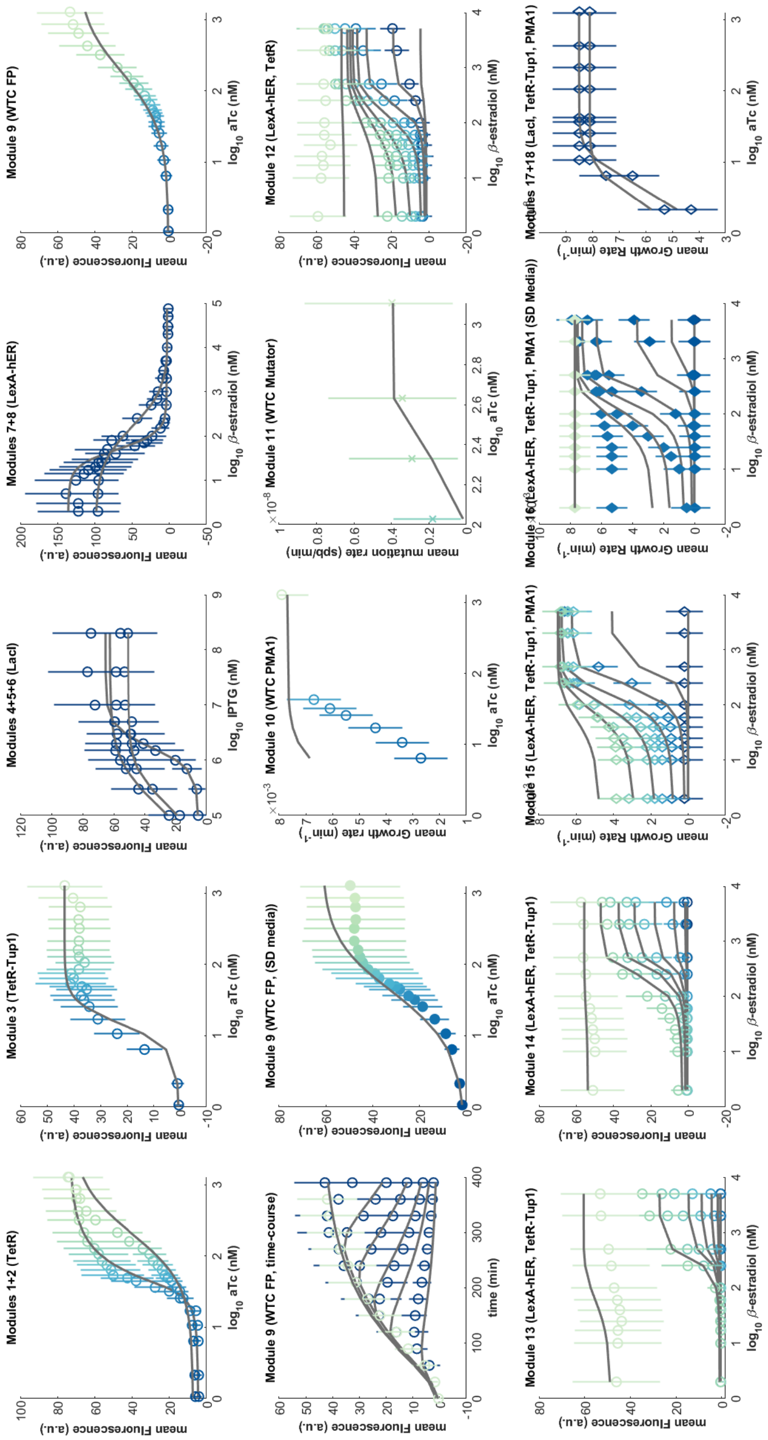


**Supplementary Figure 9: Small informative synthetic networks (modules) measured to infer the parameter values.**

The strains/modules are listed in Supplementary Table 2A. Fluorescence output (circles) was measured by flow cytometry and normalized by cell size. Growth rates (diamonds) were estimated from growth curves obtained with the GrowthProfiler (see STAR Methods) and mutation rates (crosses) were obtained by performing a fluctuation assay (see STAR Methods and Supplementary Figure 8C&D). Symbols show experimental means +/- standard deviation and lines are simulations with the parameter set that showed the best fit to the experimental data (see Supplementary Table 2D). Colors refer to different aTc concentrations: dark blue means no or a small aTc dose and a more green color shows a higher dose of aTc (up to 600 ng/mL). Modules that were measured in SD media instead of YPD media show filled data points.

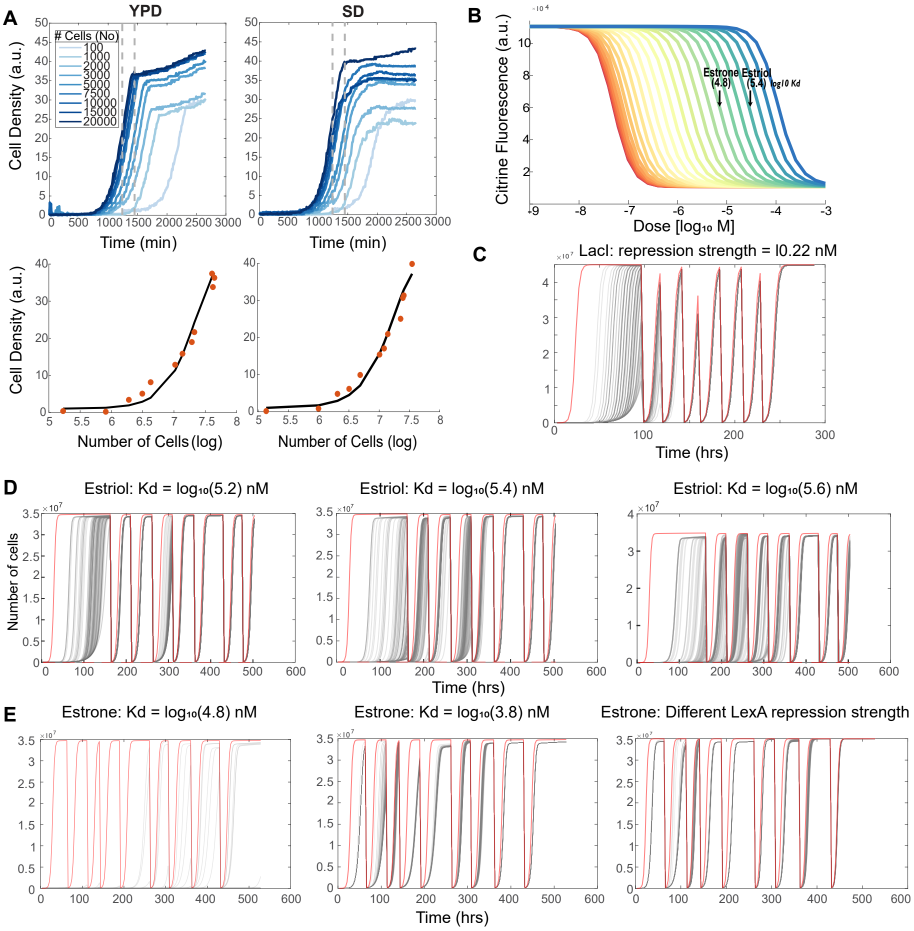


**Supplementary Figure 10: Using a parametrized model to capture the evolution trajectories.**
**(A)** Growth curves for nine different cultures, each inoculated with a different number of cells, in both YPD (left) and SD (right) media. At two different time points (grey lines), the number of cells in each culture was analyzed by CoulterCounter. The measured number of cells was plotted against the corresponding cell density (orange dots). A curve (black line) was fitted through the experimental data and subsequently used in the model to provide model outputs in cell density (a.u.). **(B)** Simulated dose-response curves for different receptor affinities. Black arrows: simulated dose-response curves matching the experimental curves for estrone and estriol (based on calculated EC50, shown in Supplementary Figure 1A). (**C)** Growth curves (in numbers of cells) obtained from simulating the LacI evolution experiment (that were subsequently used to calculate the mid-points). **(D)** Simulated growth curves for the estimated starting affinity from (B) (middle) and for slightly different starting affinities for the estriol evolution experiments. **(E)** Simulated growth curves for the estimated starting affinity from (B) (left) and for two alternative scenarios for the estrone evolution experiments: (i) a beneficial LexA-

hER mutation happened already in the diversity culture (middle) or (ii) an early mutation in LexA increased its repression strength (right).

**A**

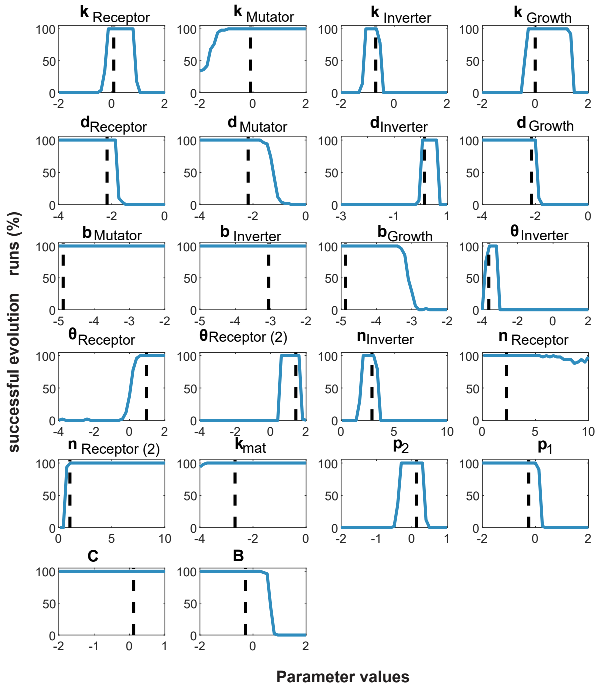


**B**

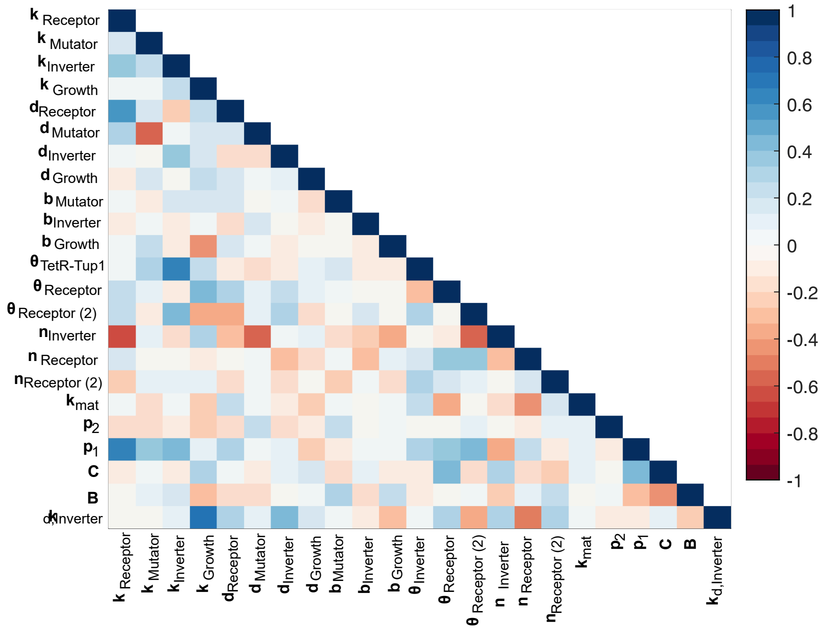


**Supplementary Figure 11: Influence of parameter sensitivities and correlations on the Evolverator model and design.**

**(A)** Success rate of evolution experiments upon changing each model parameter in isolation. Parameter meanings are explained in STAR Methods and Supplementary Table 2C. The black line in each panel indicates the parameter value obtained from parameter estimation on small, experimental modules (Supplementary Table 2D). **(B)** Heatmap depicting the pairwise-correlation between parameters of the parameterized Evolverator model. Correlations were obtained by sampling (see STAR methods).
